# Distal conformational steering by N-terminal pyroglutamylation enables subtype-selective GPCR activation across *Aplysia* PRXamide and human Neuromedin U signaling

**DOI:** 10.64898/2026.07.28.741146

**Authors:** Jian-Hui Chang, Wei-Jia Liu, Shao-Qian Wu, Elena V. Romanova, Cheng-Yi Liu, Cui-Ping Liu, Xing Pan, Fan Li, Xue-Ying Ding, Rui-Ting Mao, Hui-Ying Wang, Ju-Ping Xu, Ping Fu, Yi-Long Zhang, Qing-Chun Jin, Yan-Chu-Fei Zhang, Guo Zhang, Jonathan V. Sweedler, Jian Jing

## Abstract

Post-translational modifications (PTMs) diversify neuropeptide function, yet how minimal modifications encode receptor specificity without direct contact remains a fundamental challenge in chemical biology. Pyroglutamylation (pQ), a prevalent N-terminal PTM in bioactive peptides, introduces a rigid cyclic constraint, yet its mechanistic role in receptor signaling is unclear. Here, using two newly identified endogenous Aplysia PRXamide receptors (ApPRXa R1 and ApPRXa-R2) as a model system, we find that the same ligand, MMG2-pDPb (pQPPLPRYamide), produces opposite functional outcomes: pQ suppresses ApPRXa-R1 activation while enhancing ApPRXa-R2 activation. In vitro and in silico analyses demonstrate that the N-terminal pQ/Q remains solvent-exposed and does not directly contact receptor residues. Instead, pQ reshapes the ligand conformational ensemble and redistributes interaction networks across shared receptor contact sites. Strikingly, this molecular logic extends to mammalian Neuromedin U (NmU) receptors, as canine NmU (pQFLFRPRNamide) similarly biases subtype preference of human NmU receptors. Both static and dynamic analyses further reveal that receptor pocket mechanics determine the direction of this modulation: a loose and permissive pocket better accommodates the rigid pQ-constrained ligand, whereas a more compact pocket favors the non-pyroglutamylated ligand. These findings define a “PTM distal steering” mechanism that bridges “lock-key” and “induced-fit” paradigms and establish a general principle by which a minimal, non-contacting modification encodes receptor preference through ligand conformational biasing and pocket-dependent permissiveness, providing a chemical framework for optimizing stable, conformation-biased neuropeptide analogs.

## Introduction

Post-translational modifications (PTMs) expand the functional and chemical diversity of neuropeptides, many of which rely on such modifications for efficient activation of G protein-coupled receptors (GPCRs) (1–4), thereby regulating diverse physiological processes (5–13). Canonically, PTMs are thought to modulate neuropeptide activity through direct ligand-receptor interactions (14–19), enhancing receptor affinity or signaling efficacy. However, the mechanistic basis by which certain PTMs, such as the N-terminal pyroglutamylation (conversion to pyroglutamic acid, pQ), can exert subtype-selective modulation without directly engaging receptor residues remain unclear. Here, we investigate how this N-terminal PTM may reconfigure ligand-receptor interaction dynamics via distal conformational control.

pQ introduces a rigid five-membered lactam ring and is prevalent among neuropeptides (20–23). It is best known for conferring resistance to aminopeptidase degradation, as observed in GnRH, TRH and Tachykinin (22, 24, 25). Beyond this stabilization role, pQ has been proposed to influence receptor signaling through structural effects rather than forming direct ligand-receptor contacts (20, 26–31). For example, pQ may stabilize preferred ligand configurations (26, 28), potentially through restriction of conformational flexibility as demonstrated in solution-based studies (32, 33), or maintenance of an optimal binding orientation to enhance GPCR affinity, as inferred from studies where its substitution reversed binding pose (30, 34). Despite these proposals, no neuropeptide system involving pyroglutamylation has yet provided direct molecular-level evidence demonstrating how such conformational effects might dictate ligand-receptor interactions and signaling outcomes. In most cases, pQ either enhances receptor activation (23–26, 28) or has negligible impact upon removal (27, 29, 31), complicating efforts to disentangle its structural or mechanistic roles from stability-related effects.

This limitation is overcome by the *Aplysia* PRXamide model system. The PRXamide families (referred to as the Neuromedin U (NmU) family in vertebrates), defined by a C-terminal PRXamide motif, is conserved across protostomes and deuterostomes (35–39). Here, we described two newly identified *Aplysia* PRXamide receptors, ApPRXa-R1 and ApPRXa-R2, and discovered that they respond oppositely to the same pQ-modified ligand, MMG2-pDPb (pQPPLPRYamide) (40). ApPRXa-R2 follows the canonical pattern in which pQ enhances ligand potency, similar to many other pQ-containing peptides (24–26, 28) including *Aplysia* Tachykinins (23). In striking contrast, ApPRXa-R1 exhibits reduced responsiveness upon pQ modification. This bidirectional, subtype-specific effect of pQ is unprecedented in neuropeptide signaling. More importantly, such opposite regulation suggests that the outcome is determined not solely by the presence of pQ, but by receptor subtype-specific properties, such as distinct interaction networks and/or differences in binding-pocket environments.

In the *Aplysia* PRXamide system, an integrated strategy combining AI-assisted peptide-GPCR tools (41, 42) and residue mutagenesis reveals that N-terminal pQ remains solvent-exposed and does not directly contact receptor residues. Instead, its lactam ring constrains the peptide conformational ensemble, reweighting interaction strengths across a largely shared set of receptor contacts. Extending this molecular logic to mammals, canine endogenous pQ-NmU-7 (pQFLFRPRNamide) (43) biases activation of human NmU receptor subtypes through an analogous non-contacting mechanism. Both static pocket and molecular dynamics analyses based on recent high-resolution cryogenic electron microscopy (cryo-EM) structures of human NmU receptors (17, 18) further show that the direction of this distal modulation is determined by a key molecular variable that is largely overlooked in the prevalent “induced-fit” framework (44–48): the intrinsic properties of receptor orthosteric binding pocket. Specifically, the more permissive and looser hNmU-R1 pocket better accommodates the rigid pQ-NmU-7, whereas the more compact hNmU-R2 pocket preferentially favors the flexible non-pyroglutamylated NmU-8, which may also explain the bidirectional pQ regulation observed in the *Aplysia* PRXamide system.

Overall, our work reveals an indirect “PTM distal steering” mechanism in neuropeptide-GPCR signaling, where a non-contacting N-terminal lactam governs receptor activation through conformational biasing and pocket-dependent accommodation. This mechanism refines classical “lock-key” (49) and “induced-fit” (44) models by introducing an intermediate mode of molecular recognition. In this mode, ligand-borne conformational constraints are differently interpreted by receptor pockets with distinct permissiveness. More broadly, this work establishes pyroglutamylation as a versatile N-terminal chemical modification for tuning peptide-GPCR recognition and provides a rational design principle for optimizing stable, conformation-biased neuropeptide analogs.

## Results

### Establishment of an endogenous PRXamide system for probing N-terminal pyroglutamylation

To establish a native PRXamide system for probing the role of N-terminal pyroglutamylation (pQ), we first employed liquid chromatography-mass spectrometry (LC-MS/MS) (50, 51) to confirm the endogenous coexistence of MMG2-DPb and its pyroglutamylated counterpart MMG2-pDPb in *Aplysia* central ganglia **(Figs. S1, S2 and Table S1).** In parallel, bioinformatics and phylogenetic analyses identified four candidate PRXamide/Neuromedin U-like Class A GPCRs as their potential cognate receptors **(Figs. S3-S5 and Table S2)**. Broad functional screening against a panel of all known *Aplysia* PRXamide-related peptides, including MMG2-DPs, pleurins, SCPs and D-type pleurins, deorphanized ApPRXa-R1 and ApPRXa-R2 as cognate PRXamide receptors **(Fig. 1A and Figs. S6-S8)**. D-type pleurins did not activate any of the candidate receptors under these conditions, and therefore were not tested further. Detailed identification and screening procedures are provided in the **Supporting Results and Discussion**.

**Figure 1.**
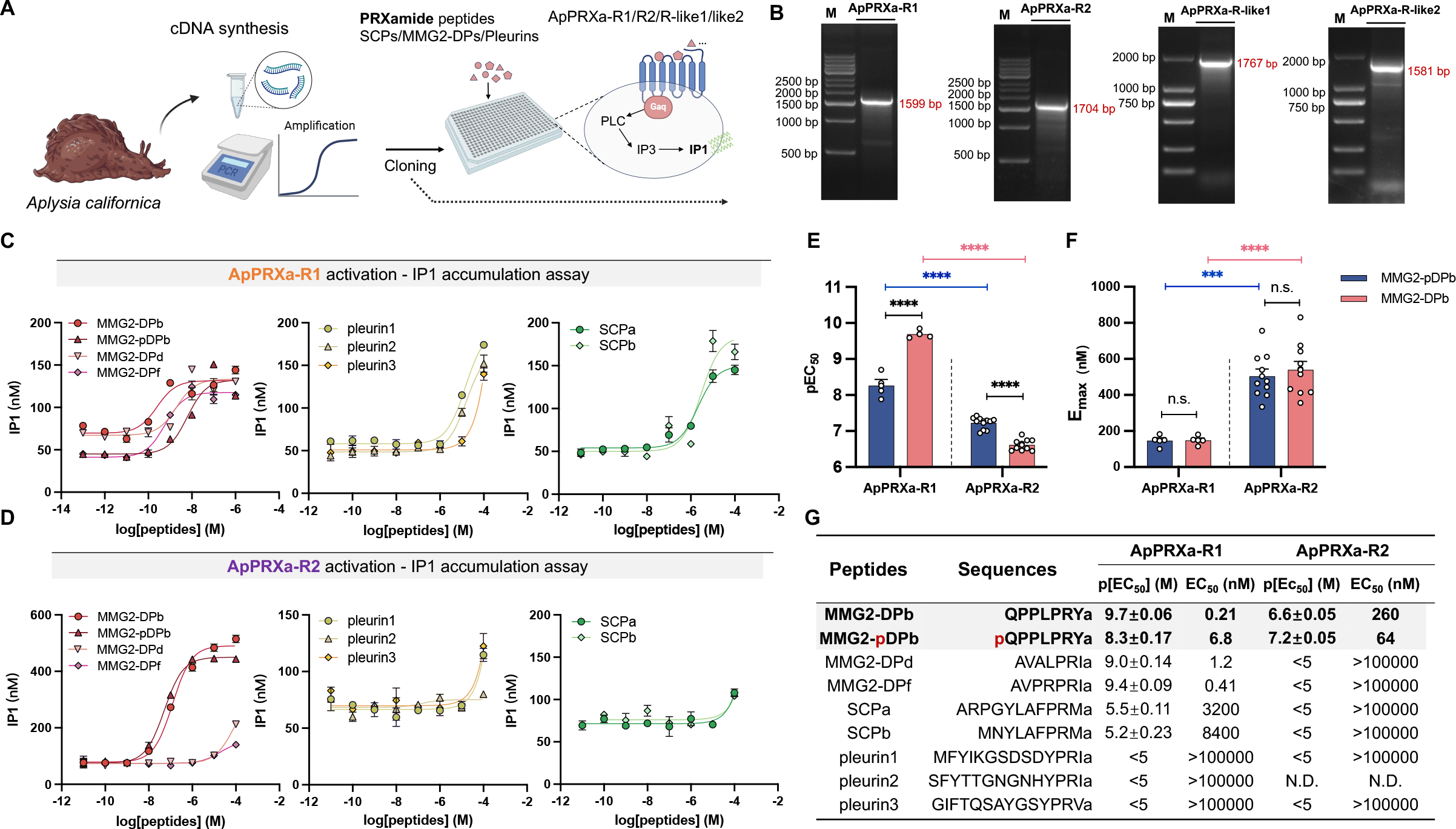
Identification of ApPRXamide receptors. **A**, Cloning of four putative ApPRXamide receptors and detection of the activation profiles induced by three classes of ApPRXamide peptides, as measured by the IP1 accumulation assay. **B,** The PCR products of putative ApPRXamide receptors (left to right): ApPRXa-R1 with a length of 1,599 bp, ApPRXa-R2 with a length of 1,704 bp, ApPRXa-R-like1 with a length of 1,767 bp, ApPRXa-R-like2 with a length of 1,581 bp. Lane 1: DNA marker (M); Lane 2: the target gene. **C-D,** Representative examples of dose-response curves showing activation of ApPRXa-R1 (**C**) and ApPRXa-R2 (**D**) in CHO-K1 cells by MMG2-DPs, pleurins and SCPs. **E,** Comparison of p[EC_50_] values for ApPRXa-R1 and ApPRXa-R2 activation by MMG2-pDPb and MMG2-DPb shown in (**C-D**), n ≥ 3. Two-way ANOVA, F_peptide_ (1, 25) = 27.41, p < 0.0001, F_receptor_ (1, 25) = 695.1, p < 0.0001. **F,** Comparison of E_max_ of ApPRXa-R1/R2, n ≥ 3. Two-way ANOVA, F_peptide_ (1, 24) = 0.1468, p > 0.05, F_receptor_ (1, 24) = 56.76, p < 0.0001. Bonferroni post-hoc test: n.s., not significant; **P < 0.01; ***P < 0.001; ****P < 0.0001. Error bar: SEM. **G,** Summary of the average p[EC_50_] and EC_50_ values shown in (**C-D**), with peptide sequences listed. IP1, inositol monophosphate; CHO-K1, Chinese hamster ovary-K1; p[EC_50_], -log_10_(EC_50_); E_max_, the maximum IP1 concentration.

### A single N-terminal lactam enables bidirectional GPCR subtype tuning

We next generated full concentration-response curves to quantify ligand potency at ApPRXa-R1 and ApPRXa-R2 **(Fig. 1 C, D)**. ApPRXa-R1 showed broad responsiveness across ApPRXamide classes, with the highest potency toward MMG2-DPs (EC_50_ = 0.21 nM) **(Fig. 1 C, G)**. In contrast, ApPRXa-R2 displayed more selective activation, responding strongly only to MMG2-DPb and MMG2-pDPb, with EC_50_ values of 260 nM and 64 nM, respectively **(Fig. 1 D, G)**. Together, these data identify ApPRXa-R1 and ApPRXa-R2 as functional receptors for MMG2-DPs, with additional receptor analyses provided in the **Supporting Results and Discussion and Figs. S9-S12**. The two receptors also differed in maximal efficacy (E_max_), with ApPRXa-R2 exhibiting higher E_max_ values than ApPRXa-R1 despite lower apparent potency **(Fig. 1F)**, possibly reflecting difference in receptor regulation (52) or desensitization kinetics (23, 53–55).

Strikingly, comparison of MMG2-DPb with its pyroglutamylated counterpart MMG2-pDPb revealed opposite effects of the same N-terminal pQ. At ApPRXa-R2, pQ enhanced ligand potency, consistent with its commonly observed potentiating role (24–26, 28). At ApPRXa-R1, however, pQ reduced ligand potency relative to MMG2-DPb **(Fig. 1 E, G)**. Thus, a single N-terminal lactam modification converts a minimal chemical difference into bidirectional GPCR subtype outputs. More importantly, such a bidirectional activation pattern indicates that pQ does not function as a universal stabilizing or affinity-enhancing group, but instead acts in a receptor-dependent manner, suggesting that the outcome must depend on subtype-specific interaction networks and possibly on how receptor-specific binding pockets accommodate the pQ-imposed ligand conformation.

### Docking reveals a non-contacting role of N-terminal pQ in reweighting receptor-contact networks

To obtain the structural basis of this bidirectional regulation, we modeled ApPRXa-R1 and ApPRXa-R2, and docked MMG2-pDPb and MMG2-DPb into the receptor binding pockets using HPEPDOCK (56), with model quality evaluation and diffdock (57) cross-validation provided in **Figs. S13-S17 and Supporting Results and Discussion**.

In all four ligand-receptor complexes, the C-terminal PRXamide pharmacophore inserted into the predicted orthosteric pockets (58), whereas the N-terminal Q or pQ remained solvent-exposed. The overall conformations thus point to the C-terminal PRXamide motif as the principal active determinant involved in direct contact with the receptors, while the peptide’s extended, linear backbone effectively projects the N-terminus outside the pocket, so on purely positional grounds, the N-terminal pQ/Q is unlikely to form direct contacts with receptor residues.

Despite the absence of direct pQ/Q-receptor interactions, MMG2-pDPb and MMG2-DPb adopted distinct conformations within both ApPRXa-Rs **(Fig. 2 A, B, Fig. 3 A, B and Fig. S18 A, B)**. Mapping of receptor-contact residues showed strong overlap with experimentally defined interaction networks in human homologous Neuromedin U receptors (17, 18), supporting the relevance of the modeled binding interface **(Figs. S19, S20 and Table S3)**. Notably, MMG2-pDPb and MMG2-DPb engaged largely shared receptor-contact sites, but differed in the strength and geometry of interactions formed by central and C-terminal residues **(Fig. 2 C, D, Fig. 3 C, D, and Table S4)**. These observations suggest that the N-terminal pQ may act as a distal conformational constraint that reweights activation-related interactions at core sites and ultimately affects receptor activity, providing an initial structural basis for its opposite effects on ApPRXa-R1 and ApPRXa-R2.

**Figure 2.**
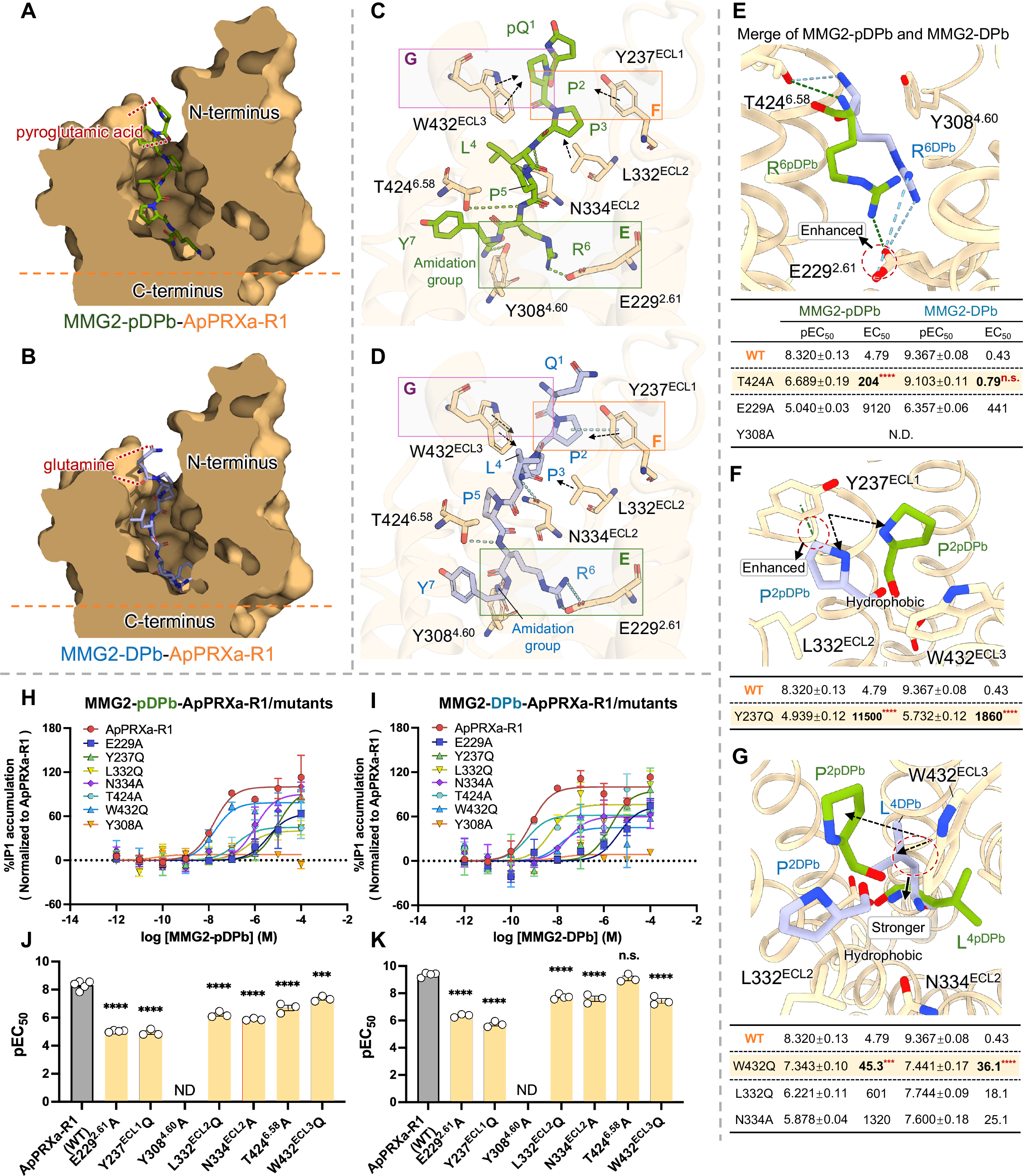
Binding modes of MMG2-pDPb and MMG2-DPb in ApPRXa-R1 from molecular docking and receptor mutagenesis. **A-B**, The cutting face of the ligand binding pocket in the (**A**) MMG2-pDPb-and (**B**) MMG2-DPb-bound ApPRXa-R1 structure. MMG2-pDPb and MMG2-DPb are represented as green and blue sticks respectively, with their N-terminal pyroglutamic acid and glutamine marked by red lines. ApPRXa-R1 is depicted as light orange surfaces. **C-D**, Overall interactions between MMG2-pDPb (**C**) and MMG2-DPb (**D**) with ApPRXa-R1. ApPRXa-R1 is shown as light orange cartoons with its residues represented as orange sticks. Hydrogen bonds and salt bridges are indicated by (**C**) green and (**D**) blue dashed lines. All receptor binding sites and corresponding ligand residues are labeled. **E-G,** Comparison of detailed interactions between MMG2-pDPb/DPb and ApPRXa-R1. The merged maps of the three regions are indicated in different colored panels in (**C-D**). Differences in interactions are observed for residues Arg^6^ (**E**), Pro^2^ (**F**) and Leu^4^ (**G**). Hydrophobic interactions are shown as dash lines with arrows, with interaction differences highlighted by red dashed circles. **H-I**, Representative examples of dose-response curves showing the activation of ApPRXa-R1 and its mutants (based on predicted binding sites) by (**H**) MMG2-pDPb and (**I**) MMG2-DPb in CHO-K1 cells, as measured by the IP1 accumulation assay. Dose-response curves are normalized to 100% of the maximal response elicited by the ligands in ApPRXa-R1. **J-K**, Summary of mutant effects on ligand potency, wildtype ApPRXa-R1 was used as a control. Panel (**J**) corresponds to (**H**). n ≥ 3. One-way ANOVA, F (6, 17) = 115.9, p < 0.0001. Panel (**K**) corresponds to (**I**). n ≥ 3. One-way ANOVA, F (6, 16) = 107.8, p < 0.0001. Bonferroni post-hoc test: n.s., not significant; ***P < 0.001; ****P < 0.0001. Error bar: S.E.M.

**Figure 3.**
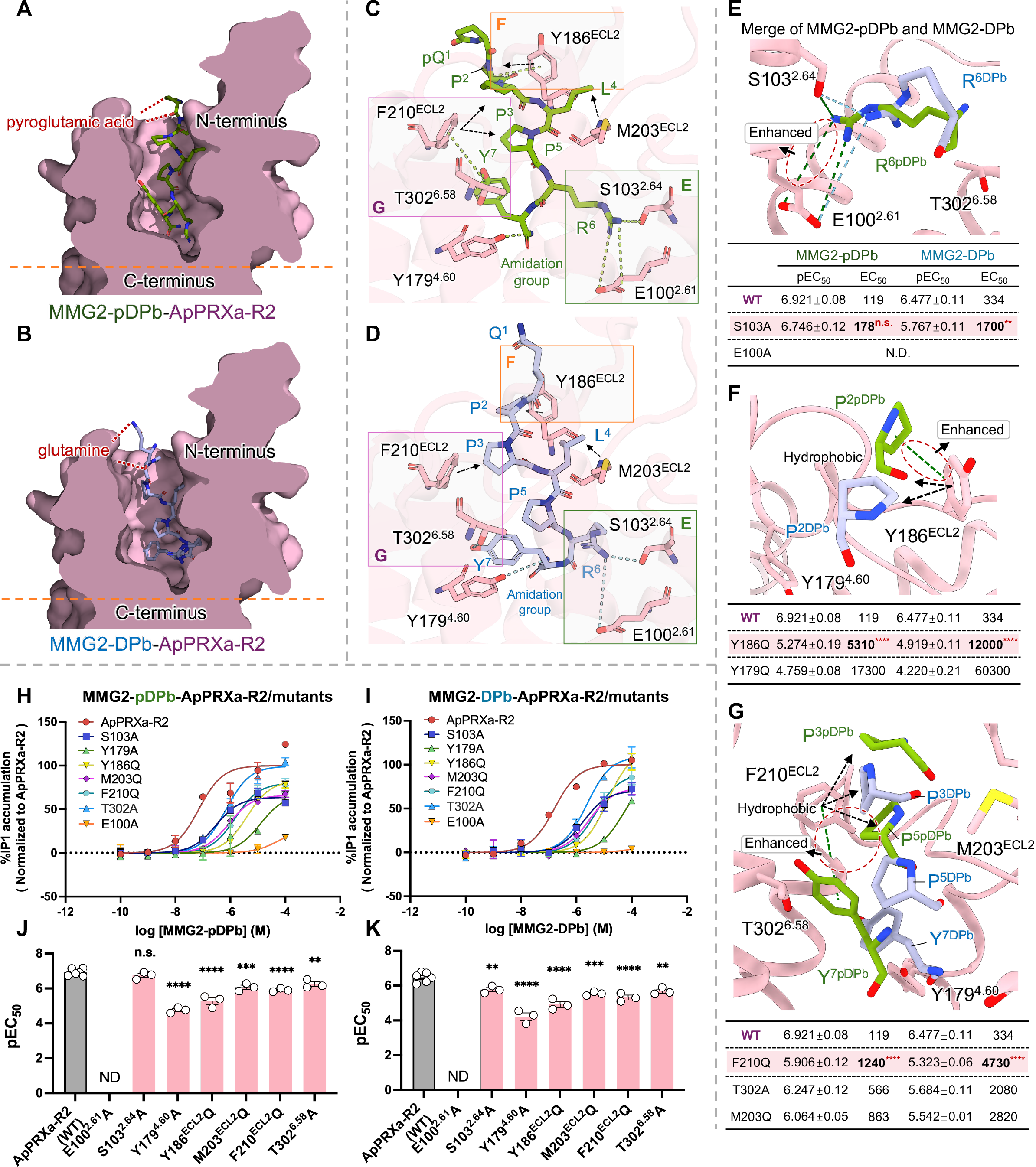
Binding modes of MMG2-pDPb and MMG2-DPb in ApPRXa-R2 from molecular docking and receptor mutagenesis. **A-B**, The cutting face of the ligand binding pocket in the (**A**) MMG2-pDPb-and (**B**) MMG2-DPb-bound ApPRXa-R2 structure. MMG2-pDPb and MMG2-DPb are represented as green and blue sticks respectively, with their N-terminal pyroglutamic acid and glutamine marked by red lines. ApPRXa-R2 is depicted as light pink surfaces. **C-D**, Overall interactions between MMG2-pDPb (**C**) and MMG2-DPb (**D**) with ApPRXa-R2. ApPRXa-R2 is shown as light pink cartoons with its residues represented as pink sticks. Hydrogen bonds and salt bridges are indicated by (**C**) green and (**D**) blue dashed lines. All receptor binding sites and corresponding ligand residues are labeled. **E-G**, Comparison of detailed interactions between MMG2-pDPb/DPb and ApPRXa-R2. The merged maps of the three regions are indicated in different colored panels in (**C-D**). Differences in interactions are observed for residues Arg^6^ (**E**), Pro^2^ (**F**), Pro^3^, Pro^5^, and Tyr^7^ (**G**). Hydrophobic interactions are shown as dash lines with arrows, with interaction differences highlighted by red dashed circles. **H-I**, Representative examples of dose-response curves showing the activation of ApPRXa-R2 and its mutants (based on predicted binding sites) by (**H**) MMG2-pDPb and (**I**) MMG2-DPb in CHO-K1 cells, as measured by the IP1 accumulation assay. Dose-response curves are normalized to 100% of the maximal response elicited by the ligands in ApPRXa-R2. **J-K**, Summary of mutant effects on ligand potency, wildtype ApPRXa-R2 was used as a control. Panel (**J**) corresponds to (**H**). n ≥ 3. One-way ANOVA, F (6, 17) = 46.46, p < 0.0001. Panel (**K**) corresponds to (**I**). n ≥ 3. One-way ANOVA, F (7, 17) = 29.82, p < 0.0001. Bonferroni post-hoc test: n.s., not significant; **P < 0.01; ***P < 0.001; ****P < 0.0001. Error bar: SEM.

### Receptor mutagenesis validates N-terminal pQ-dependent reweighting of interaction networks

To experimentally test this model, we performed site-directed mutagenesis of receptor residues predicted to interact with MMG2-DPb or MMG2-pDPb. Hydrophobic residues were generally substituted with glutamine (Q) to disrupt nonpolar contacts, whereas other residues were substituted with alanine (A). In ApPRXa-R1, mutants E229^2.61^A, Y237^ECL1^Q, L332^ECL2^Q, N334^ECL2^A, and W432^ECL3^Q significantly impaired receptor activity, while Y308^4.60^A abolished activation. In ApPRXa-R2, Y179^4.60^A, Y186^ECL2^Q, M203^ECL2^Q, F210^ECL2^Q, and T302^6.58^A markedly reduced activity, whereas E100^2.61^A led to complete inactivation **(Table S4)**.

For ApPRXa-R1, docking predicted three ligand interaction regions with notable differences between MMG2-pDPb and MMG2-DPb **(Fig. 2 E-G and Fig. 4)**. The most critical site was W432^ECL3^, which engaged in hydrophobic interactions with P^2^ of MMG2-pDPb but with L^4^ of MMG2-DPb **(Fig. 2 G)**. Given that leucine is more hydrophobic than proline (59), W432^ECL3^Q was expected to preferentially disrupt MMG2-DPb recognition. Consistent with this prediction, W432^ECL3^Q caused a larger potency loss for MMG2-DPb, even eliminating the potency difference between MMG2-pDPb and MMG2-DPb (EC_50_ = 45.5 and 36.1 nM, respectively) **(Fig. 2 G, J, K)**. These data demonstrate that W432^ECL3^-mediated hydrophobic packing is central to the N-terminal pQ-driven conformational disfavoring in receptor activation of ApPRXa-R1. Additionally, R^6^ of both ligands formed hydrogen bonds with T424^6.58^ and salt bridges with E229^2.61^. Docking suggested that the R^6^-E229^2.61^ salt bridge was weaker for MMG2-pDPb than for MMG2-DPb **(Fig. 2 E)**, making MMG2-pDPb more dependent on the T424^6.58^-R^6^ hydrogen bond. In agreement with this model, T424^6.58^A sharply decreased the potency of MMG2-pDPb (EC_50:_ from 4.79 to 204 nM), whereas the potency of MMG2-DPb remained largely unaffected (EC_50_: from 0.43 to 0.79 nM) **(Fig. 2 E, J, K)**. Furthermore, Y237^ECL1^ formed hydrophobic contacts with P^2^ of both ligands, with an additional CH-pi interaction unique to MMG2-DPb **(Fig. 2 F)**, theoretically enhancing its potency. We anticipated a relatively smaller impact of Y237^ECL1^Q on MMG2-pDPb compared to MMG2-DPb; however, Y237^ECL1^Q significantly increased EC_50_ values for both ligands **(Fig. 2 F, J, K)** without clear differentiation. These findings suggest that hydrophobic interactions mediated by Y237^ECL1^ critically influence receptor activation for both ligands, regardless of the presence of the additional CH-pi interaction, providing valuable clues for subsequent ligand-focused studies (see the next section).

**Figure 4.**
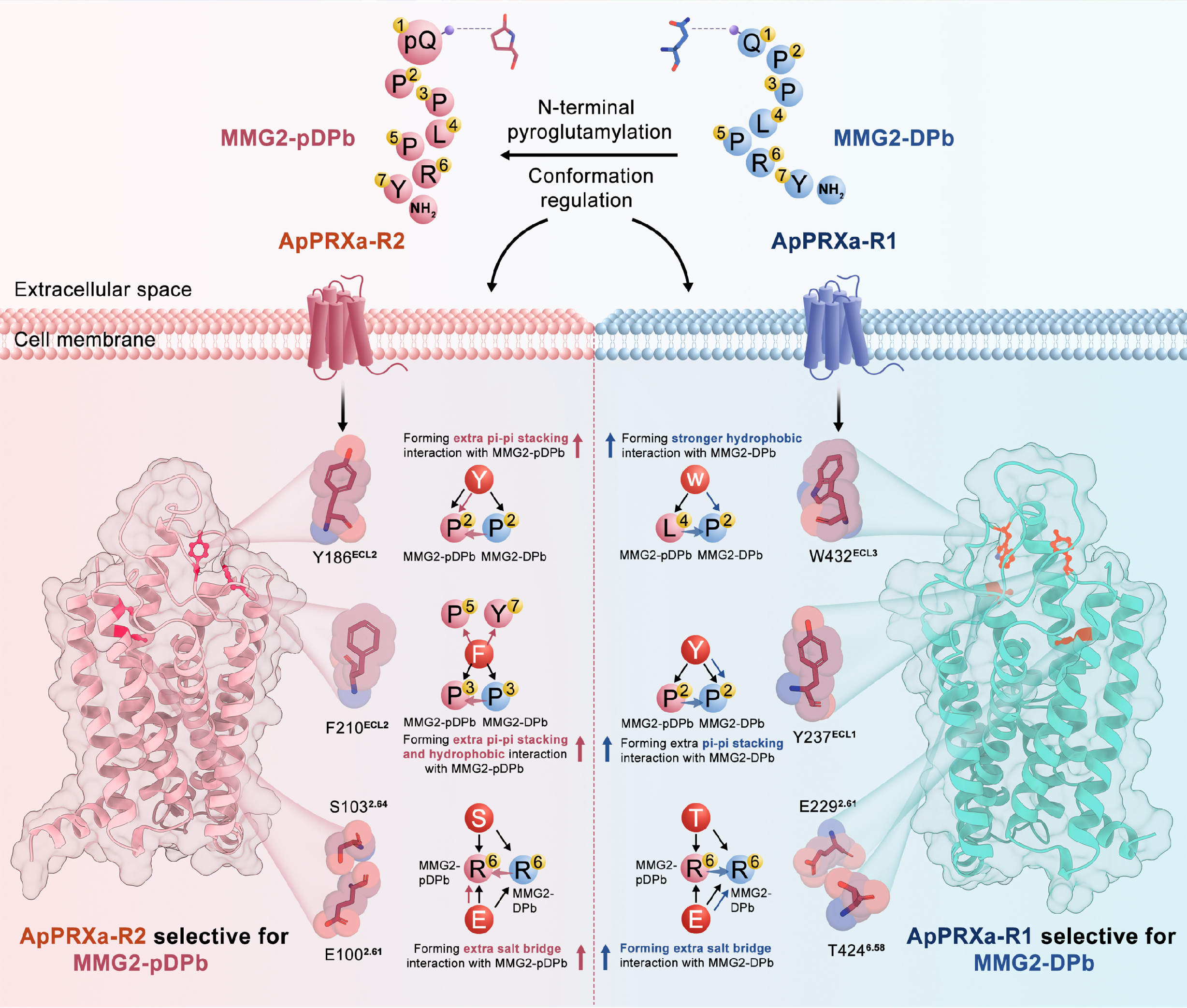
Schematic illustrating how N-terminal pyroglutamylation (pQ) remotely modulates receptor subtype-specific activation without direct contacts. In both ApPRXa-R1 and ApPRXa-R2, MMG2-pDPb (pQPPLPRYamide), which exhibits reduced conformational flexibility due to pQ, does not engage new receptor contacts but instead adopts a distinct bound conformation, resulting in differences in the number or strength of bonds with largely shared sets of interacting receptor residues. Compared with a more flexible MMG2-DPb, in ApPRXa-R2 (left, red), which may possess a loose binding pocket, the more rigid ligand with pQ strengthens key interactions with Y186^ECL2^, F210^ECL2^, S103^2.64^ and E100^2.61^, thereby potentiating receptor activation. In contrast, in ApPRXa-R1 (right, cyan), whose pocket is intrinsically compact, the same rigid pQ-modified ligand disrupts the interaction network formed by the more flexible MMG2-DPb, including contacts involving W432^ECL3^, Y237^ECL1^, E229^2.61^ and T424^6.58^, resulting in reduced receptor activity. This bidirectional, subtype-specific mechanism—enhancement in ApPRXa-R2 versus inhibition in ApPRXa-R1—accounts for the preferential activation of ApPRXa-R2 by MMG2-pDPb and of ApPRXa-R1 by the non-cyclized form MMG2-DPb.

For ApPRXa-R2, three interaction regions modulated by pQ were also identified **(Fig. 3 E-G and Fig. 4)**. R^6^ of both ligands interacted with S103^2.64^ and E100^2.61^ via hydrogen bonds and salt bridges, with the salt bridge being stronger for MMG2-pDPb **(Fig. 3 E)**. As expected, S103^2.64^A slightly affected MMG2-pDPb potency (EC_50_: from 119 to 178 nM), but significantly impaired MMG2-DPb activation (EC_50_: from 334 to 1700 nM) **(Fig. 3 E, J, K)**. Additionally, F210^ECL2^formed a hydrophobic network with P^3^ and P^5^ and pi-pi stacking with Y^7^ in MMG2-pDPb, whereas its interaction with MMG2-DPb was limited mainly to P^3^ **(Fig. 3 G)**. Despite predicting a more pronounced effect on MMG2-pDPb, F210^ECL2^Q comparatively increased EC_50_ for both ligands **(Fig. 3 G, J, K)**, supporting that the conserved hydrophobic interactions involved in F210^ECL2^ and P^3^ play an indispensable role in maintaining the receptor binding interface. Similarly, Y186^ECL2^ formed dual hydrophobic and CH-pi contacts with P^2^ of MMG2-pDPb but only hydrophobic contacts with MMG2-DPb **(Fig. 3 F)**. Although Y186^ECL2^Q had no selective effects on the two ligands **(Fig. 3 F, J, K)**, the strong loss of potency for both ligands supports its role in stabilizing the ApPRXa-R2 binding pocket.

Beyond individual contacts, the two receptors differ in overall pocket architecture. In ApPRXa-R2, MMG2-pDPb is cradled within a hydrophobic pocket defined by Y186^ECL2^, F210^ECL2^ and M203^ECL2^, which engaged multiple ligand positions, including P^2^, P^3^, L^4^ and P^5^ **(Fig. 3 C, Fig. S20 and Table S4)**. This distributed hydrophobic interface may provide a relatively loose and permissive environment for pQ-mediated potentiation. Consistently, mutations at S103^2.64^, M203^ECL2^ and T302^6.58^ only moderately impair receptor activation **(Fig. 3 J-K and Fig. S20 E)**. By contrast, in ApPRXa-R1, Y237^ECL1^, L332^ECL2^, N334^ECL2^ and W432^ECL3^ contact a narrow set of ligand positions, mainly P^2^ and P^3^, generating a more compact and tightly optimized recognition network **(Fig. 2 C, Fig. S20 and Table S4)**. In this compact space, individual mutations dramatically impaired receptor activation **(Fig. 2 J-K and Fig. S20 E)**, indicating reduced tolerance to ligand conformational perturbation. Structural alignments further supported that pQ amplifies subtype-specific conformational divergence between ligand-bound states **(Fig. S13 C and Fig. S18 C, D)**. These features also align with the broader ligand tolerance and generally lower EC_50_ values of ApPRXa-R1 compared with ApPRXa-R2 when activated by either MMG2-pDPb or MMG2-DPb **(Fig. 1 G and Fig. S21)**. Together, at the molecular level, pQ remodels the ligand conformation and redistributes the contribution of pre-existing receptor-contact networks **(Figs. 2-4)**. We therefore propose that the distinct pocket properties of ApPRXa-R1 and ApPRXa-R2 may differentially interpret the pQ-imposed regulation, providing a plausible basis for the observed subtype-selective activation.

Cell surface receptor expression analyses of FLAG-tagged mutants confirmed comparable expression levels to wild-type receptors **(Figs. S22-S24)**, indicating that EC_50_ differences reflect binding alterations rather than trafficking variations. Moreover, the EC_50_ values for FLAG-tagged receptors exhibited a similar trend to those of untagged receptors **(Figs. S23, S24)**.

### Activation of MMG2-pDPb analogs defines pQ as a distal conformational modulator

We tested a series of MMG2-pDPb analogs at both ApPRXa-R1 and ApPRXa-R2, and modeled their receptor-bound conformations to further define the ligand-side determinants of pQ-dependent subtype tuning. Replacing the N-terminal pQ with alanine generated [Ala^1^]MMG2-pDPb, which resembled MMG2-DPb in both docking pose, recognition sites **(Fig. 5 B, C and Table S5)** and ligand potency **(Fig. 5 D-F, J, K)** rather than MMG2-pDPb. This further supports the idea that the pQ effects arise primarily from conformational modulation rather than direct receptor contacts. Next, we examined an acetylated analog, [Ac-Ala^1^]MMG2-pDPb, to test whether N-terminal capping with charge neutralization alone could account for the pQ-dependent effects. This analog partially reproduced the inhibitory effect of MMG2-pDPb at ApPRXa-R1, but failed to fully restore MMG2-pDPb-like potency at ApPRXa-R2 **(Fig. 5 D-F, J, K)**, indicating that the pQ lactam provides a more specific structural constraint for productive ApPRXa-R2 activation.

**Figure 5.**
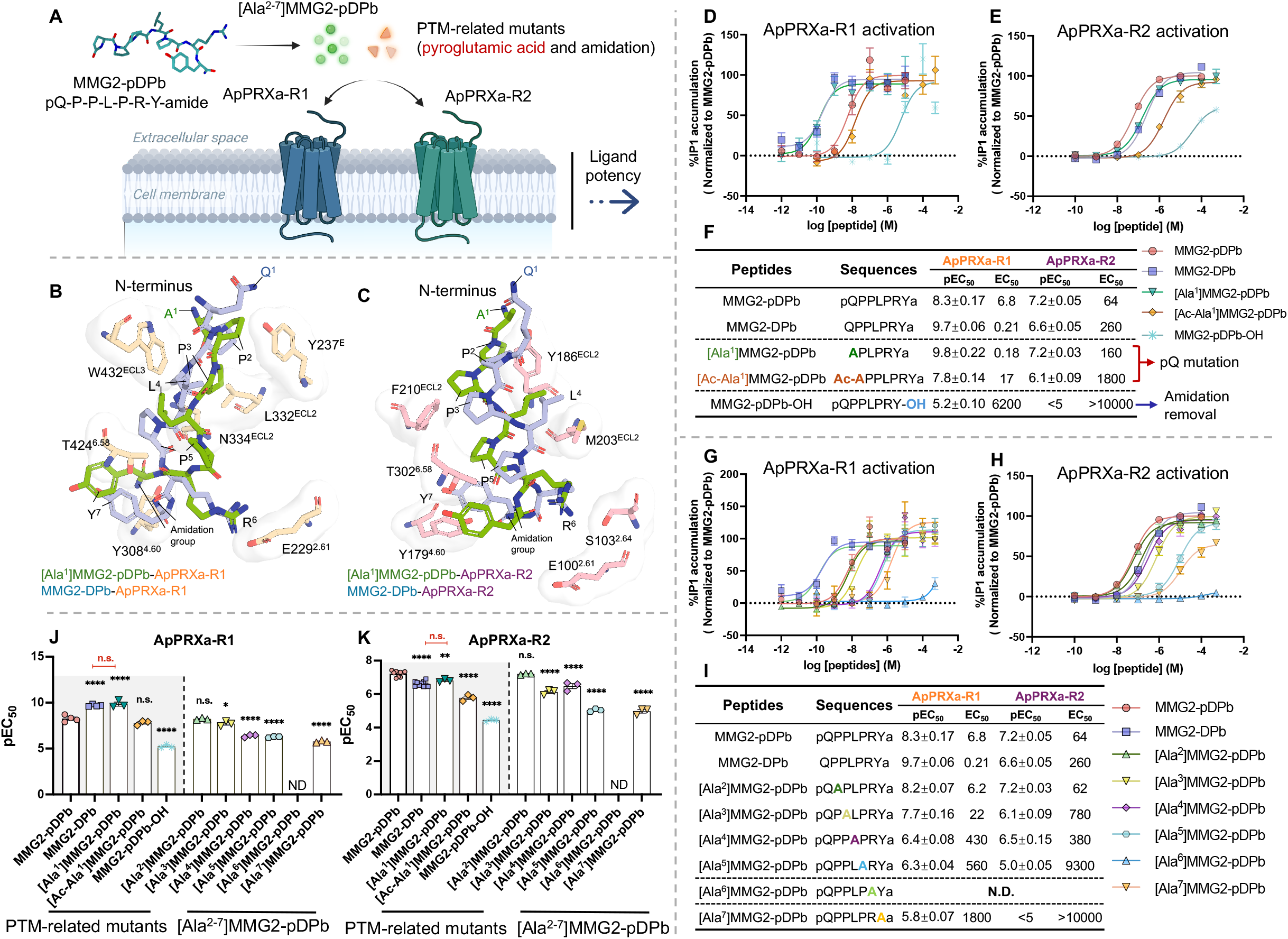
Activation of ApPRXa-Rs by MMG2-pDPb analogs. **A**, There were two types of MMG2-pDPb analogs: with one substituting the residues at positions 2-7 with Ala, and the other modifying the post-translational modifications (PTM) including pyroglutamic acid and amidation group. **B-C,** Comparison of conformations of [Ala^1^]MMG2-pDPb and MMG2-DPb in ApPRXa-R1 (**B**) and ApPRXa-R2 (**C**). [Ala^1^]MMG2-pDPb and MMG2-DPb are represented as green and blue sticks respectively. ApPRXa-Rs residues are shown as sticks with white surfaces. All receptor binding sites and corresponding ligand residues are labeled. **D-E,** Representative examples of dose-response curves showing the activation of ApPRXa-R1 (**D**) and ApPRXa-R2 (**E**) by PTM-related analogs. **F,** Summary of the average p[EC_50_] and EC_50_ values shown in (**D-E**), with peptide sequences listed. **G-H,** Representative examples of dose-response curves showing the activation of ApPRXa-R1 (**G**) and ApPRXa-R2 (**H**) by [Ala^2-7^]MMG2-pDPb. **I,** Summary of the average p[EC_50_] and EC_50_ values shown in (**G-H**), with peptide sequences listed. Dose-response curves are normalized to 100% of the maximal response elicited by MMG2-pDPb in receptors. **J-K,** Group data comparing the effects of all MMG2-pDPb analogs on ApPRXa-R1 (**J**) and ApPRXa-R2 (**K**). MMG2-pDPb was used as a control. Panel (**J**) corresponds to (**D**) and (**G**). n ≥ 3. One-way ANOVA, F (9, 22) = 153.2, p < 0.0001. Panel (**K**) corresponds to (**E**) and (**H**). n ≥ 3. One-way ANOVA, F (9, 34) = 151.3, p < 0.0001. Bonferroni post-hoc test: n.s., not significant; *P < 0.05; **P < 0.01; ****P < 0.0001. Error bar: SEM.

For C-terminal amidation, its removal (MMG2-pDPb-OH) sharply increased EC_50_ values, reaching 6.2 μM for ApPRXa-R1 and exceeding 10 μM for ApPRXa-R2, which could not be fully activated **(Fig. 5 D-F, J, K)**. Structurally, amidation loss resulted in the disappearance of hydrogen bonds and affected the conformation of R^6^ and Y^7^ **(Figs. S25, S26 and Table S5)**, highlighting its important role as a primary activation pharmacophore.

In addition to PTMs, alanine-scanning of residues 2-7 ([Ala^2^]-[Ala^7^]MMG2-pDPb) revealed a gradient of functional contributions **(Fig. 5 G-K, Figs. S25, S26 and Table S5)**. For ApPRXa-R1 system, Ala^2^ substitution merely shifted the hydrophobic interaction between Y237^ECL1^ and P^2^ to P^3^. Similarly, in ApPRXa-R2, A^2^ maintained hydrophobic contact with Y186^ECL2^. Potency of [Ala^2^]MMG2-pDPb was nearly identical to the native peptide in both receptors, indicating no critical interaction loss in either case. In contrast, [Ala^3^]MMG2-pDPb abolished the hydrophobic interaction between F210^ECL2^ and P^3^ in ApPRXa-R2, leading to a marked reduction in receptor activation, while the interaction pattern and efficacy remained largely unchanged in ApPRXa-R1. This demonstrates the dual role of the hydrophobic “anchor” interactions in peptide binding, while secondary contacts modulate fine conformational tuning. Notably, [Ala^6^]MMG2-pDPb exhibited a complete loss of activity in both receptors. Structural modeling and mutagenesis data (**Fig. 2 J, Fig. 3 J, Figs. S25, S26 and Table S5)** indicated the absence of key hydrogen bonds and salt bridges originally formed with R^6^, particularly the salt bridges involving E229/E100^2.61^, which proved essential for activation (EC_50_-E229^2.61^A: 9120; E100^2.61^A: inactive). This likely reflects the strong electrostatic attraction between the guanidino group of arginine and the carboxyl group of glutamic acid (60), which is consistent with findings from other PRXamide/Neuromedin U structure-activity relationship studies (17, 18, 61). A general trend was observed whereby alanine substitutions closer to the C-terminal motif led to a progressive loss of potency (EC_50_ fold change: 3.2-265 in ApPRXa-R1; 12-156 in ApPRXa-R2). Interestingly, [Ala^4^]MMG2-pDPb was less disruptive than [Ala^5^] MMG2-pDPb in ApPRXa-R2, possibly due to the boundary location of L^4^ and the conformational flexibility imparted by its side chain, suggesting its involvement in receptor selectivity. [Ala^7^] MMG2-pDPb partially activated ApPRXa-R2 (Emax ∼65% to wild-type), consistent with loss of T302^6.58^ hydrogen bonding or destabilized C-terminal amidation interactions.

### pQ-driven bidirectional steering logic extends to human NmU receptor subtypes

The subtype-selective regulation observed in the *Aplysia* PRXamide signaling **(Fig. 1 E and Fig. 4)** prompted us to ask whether an analogous distal steering logic might operate in mammalian NmU signaling. Although most mammalian NmUs lack an N-terminal Q, canine pQ-NmU-7 (pQFLFRPRNamide) **(Fig. S2 D)** uniquely carries an N-terminal pQ (43), raising the intriguing possibility that pQ may act as conserved molecular switch. To evaluate ligand-receptor interaction networks, we leveraged available cryo-EM structures of human receptor subtypes (hNmU-R1 and hNmU-R2) bound to NmU-8 (17, 18), and performed comparable docking analyses with NmU-8 and its two analogs: pQ-NmU-7 **(Fig. S27)** (43) and Q-NmU-7 (QFLFRPRNamide) (A similar modeling analysis of the insect RPCH signaling system with a pQ ligand is provided in **Supporting Results and Discussion and Fig. S28**).

Docking of NmU-8 recapitulated cryo-EM observations (17, 18), showing that N-terminal residue (Y) does not directly contact receptors **(Figs. S29 A, S30 A),** and that the identified interaction residues fully overlapped with experimentally resolved contacts **(Fig. S31)**. Consistent with this, the N-terminal residue (pQ, or Q) in both pQ-NmU-7 and Q-NmU-7 also remains solvent-exposed and does not directly engage receptor residues **(Figs. S29 B-C, S30 B-C)**.

Despite this lack of direct contact, pQ markedly alters interaction patterns. In hNmU-R1, the rigidity imparted by pQ (pQ-NmU-7) appears to optimize the side-chain positioning of L^3^, reinforcing hydrophobic packing against W320^ECL3^ and F334^7.31^, while R^5^ approached C219^45.50^ to form a polar contact **(Fig. S29 B, E, G)**. Conversely, in hNmU-R2, the same pQ constraint hinders the conformational rearrangements required to accommodate bulky side chains within a narrower binding pocket, causing a steric mismatch that diminishes the L^3^-centered hydrophobic network (involving A302^7.28^ and F305^7.31^) and disrupts polar contacts near R^5^ (with T203^ECL2^ and T205^ECL2^) **(Fig. S30 B, E, G)**. Thus, our analyses support pQ as an indirect conformational switch, changing the number or strength of bonds formed by a largely shared set of interacting receptor residues to bias subtype activation—enhancing it in hNmU-R1 while attenuating it in hNmU-R2.

To experimentally validate these predictions, we synthesized NmU-8, pQ-NmU-7 and Q-NmU-7 **(Fig. 6)**, and evaluated their potencies at hNmU-1 and hNmU-2. In hNmU-R1, pQ-NmU-7 significantly potentiated the receptor activation compared with both NmU-8 and Q-NmU-7, yielding approximately a 10-fold decrease in EC_50_ **(Fig. 6 A, C)**. In contrast, Q-NmU-7 had no obvious impact on hNmU-R1 activation **(Fig. 6 A, C)**. Intriguingly, in hNmU-R2, the regulatory pattern differed. pQ-NmU-7 modestly attenuated receptor activation relative to NmU-8, increasing EC_50_ values from 3.9 to 19.6 **(Fig. 6 B, C)**. A similar weakening trend was observed compared to Q-NmU-7, though not statistically significant **(Fig. 6 B, C)**. As observed for hNmU-R1, Q-NmU-7 did not significantly modulate hNmU-R2 activation **(Fig. 6 A-C)**. Thus, the overall activation patterns are consistent with our prediction.

**Figure 6.**
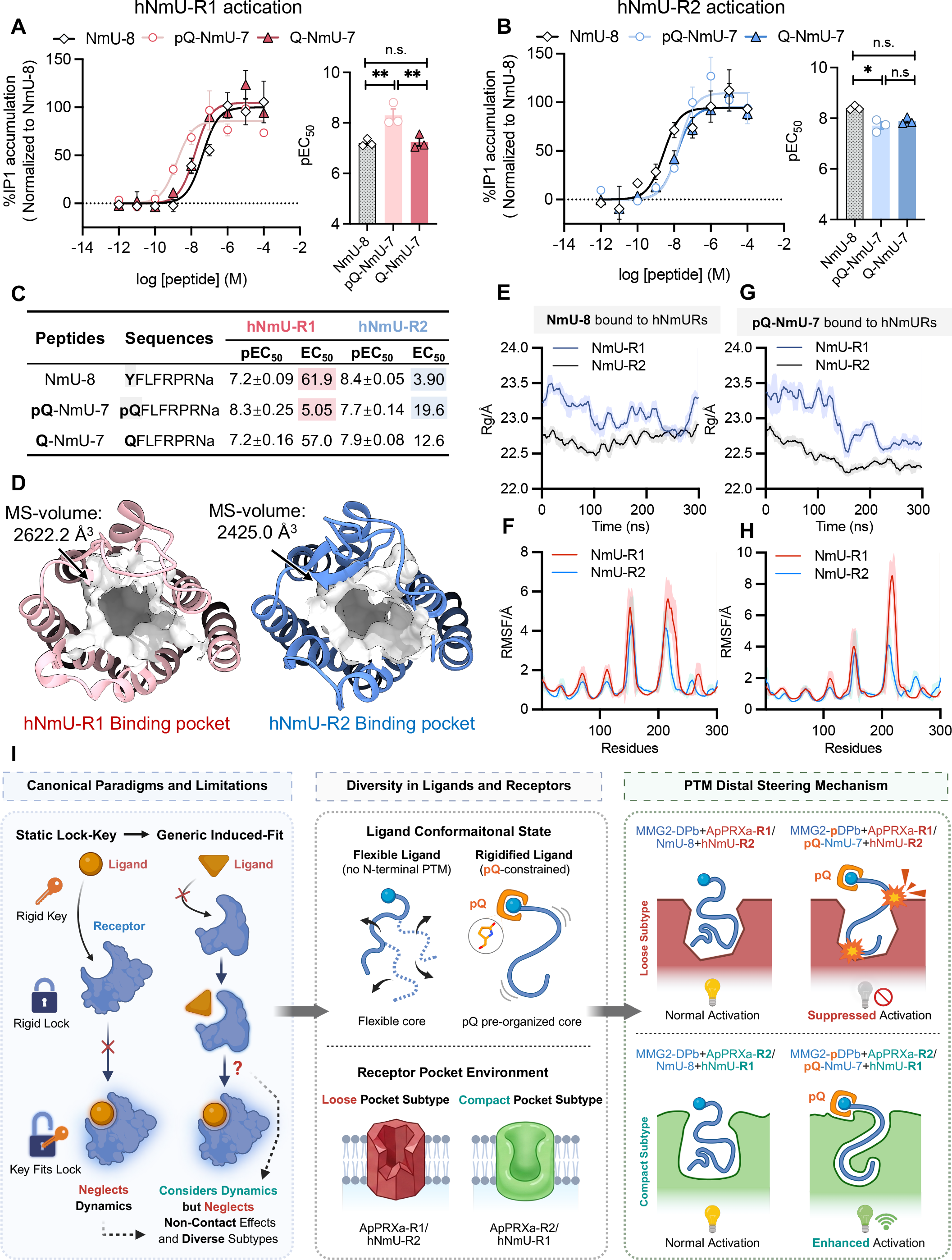
Human Neuromedin U receptor subtypes reveal pocket-dependent distal steering by N-terminal pyroglutamylation. **A-B**, Representative examples of dose-response curves and summary data showing the activation of hNmU-R1 (**A**) and hNmU-R2 (**B**) by NmU-8 (YFLFRPRNamide), pQ-NmU-7 (pQFLFRPRNamide) and Q-NmU-7 (QFLFRPRNamide). NmU-8 exhibits distinct potencies at the two receptors as previously reported (17). For assays of pQ/Q-NmU-7 on hNmU-R1, n =3. One-way ANOVA, F (2, 6) = 11.75, p < 0.01; hNmU-R2, n =3. One-way ANOVA, F (2, 6) = 13.58, p < 0.01. Bonferroni post-hoc test: n.s., not significant; *P < 0.05; **P < 0.01. Error bar: SEM. **C,** Summary of the average p[EC_50_] and EC_50_ values from (**A-B**), with peptide sequences listed. NmU-8, Neuromedin U-8; hNmU-R1/2, human Neuromedin U receptors 1/2. **D,** Orthosteric binding pockets of ligand-bound active-state hNmU-R1(red) and hNmU-R2 (blue). Receptors are shown as cartoons and CASTp-defined molecular pockets are shown as white surfaces. **E-F,** Rg (**E**) and RMSF (**F**) profiles from 300-ns MD simulations of NmU-8-bound hNmU-R1 and hNmU-R2 complexes**. G-H,** Rg (**G**) and RMSF (**H**) profiles from 300-ns MD simulations of pQ-NmU-7-bound hNmU-R1 and hNmU-R2 models. **I,** Conceptual framework for PTM distal steering mechanism in relation to classical ligand-receptor interaction models. Left panel, classical paradigms and their limitations. The lock-key model assumes rigid, pre-complementary binding and neglects conformational dynamics of both ligand and receptor. The induced-fit model incorporates receptor rearrangement but does not account for non-contacting ligand features or divergent receptor-subtype architectures. Middle panel, structural diversity of ligands and receptors illustrated by PRXamide/NmU signaling. Ligands lacking N-terminal pQ sample multiple dynamic conformations (dashed lines, arrows), whereas N-terminal pQ rigidifies the peptide and pre-organizes its core into a constrained conformation. Receptor pockets range from compact environments (red; e.g., ApRPXa-R1, hNmU-R2) to looser pockets (green; e.g., ApPRXa-R2, hNmU-R1). Right panel, PTM distal steering mechanism. Top left: a compact receptor pocket paired with a flexible, non-pQ ligand permits baseline activation. Top right: the same compact pocket challenged by a pQ-rigidified ligand induces conformational frustration and reduced activation, defining an intermediate binding mode between the lock-key and induced-fit. Bottom left: a loose pocket interacting with a flexible, unmodified ligand enables baseline activation. Bottom right: the loose pocket productively accommodates the pQ-rigidified ligand, enhancing activation via canonical induced-fit.

### Pocket permissiveness determines whether the pQ-constrained ligand is favored or penalized

The bidirectional regulation observed for canine pQ-NmU-7 further prompted us to examine whether receptor pocket properties could explain why the same N-terminal lactam constraint produced opposite effects at human NmU receptor subtypes. Previous structural study demonstrated that L^3^, F^4^ or R^5^ of NmU-8 are accommodated in a more extensive residue environment in hNmU-R1, whereas the corresponding recognition region in hNmU-R2 is more compact (17). This distinction suggested that the more conformationally rigid pQ-NmU-7 might be differentially accommodated by the binding pockets of the two receptors, thereby leading to opposite subtype outputs.

To test this hypothesis, we performed quantitative analyses using cryo-EM structures of both hNmU-R1 and hNmU-R2 (17). First, we analyzed the orthosteric pocket volumes of the ligand-bound active-state hNmU receptor structures. CASTp analysis (62) showed that the hNmU-R1 binding pocket had a larger molecular-surface (MS) volume than hNmU-R2 (2622.2 Å^3^ versus 2425.0 Å^3^) **(Fig. 6 D)**, consistent with a looser receptor environment for ligand recognition.

Second, we performed 300-ns all-atom molecular dynamics (MD) for NmU-8 bound to hNmU-R1 and hNmU-R2. Notably, radius of gyration (Rg) values were consistently higher for the hNmU-R1 complex than for the hNmU-R2 complex **(Fig. 6 E)**, suggesting a less restricted and more open overall receptor-ligand architecture. Root-mean-square fluctuation (RMSF) analysis further showed higher residue fluctuations in hNmU-R1 than in hNmU-R2, particularly around residues 200-300, including major binding regions **(Fig. 6 F)**. Parallel 300-ns MD simulations based on the pQ-NmU-7 docking models showed the same Rg and RMSF trends **(Fig. 6 G, H)**, indicating that this pocket-property difference is maintained in both NmU-8-and pQ-NmU-7-bound complexes. Root-mean-square deviation (RMSD) trajectories and free-energy landscapes (FEL) indicated that both complexes remained structurally stable during the simulations **(Fig. S32)**.

Together, these static and dynamic analyses indicated that hNmU-R1 provides a spacious and dynamically permissive pocket environment, whereas hNmU-R2 forms a more compact and restrictive architecture. This difference explains why the conformationally rigid pQ-NmU-7 is favored by hNmU-R1 but penalized by hNmU-R2, converting the same non-contacting N-terminal modification into opposite subtype-selective activation **(Fig. 6 I)**.

## Discussion

This study establishes how a minimal N-terminal chemical modification can remotely control peptide-GPCR recognition. Using the *Aplysia* PRXamide system as a molecularly tractable structure-activity relationship (SAR) model, we show that two receptor subtypes (ApPRXa-R1 and ApPRXa-R2) respond in opposite directions to the same pyroglutamylated ligand, MMG2-pDPb. This bidirectional activity cannot be explained by a simple stabilizing or affinity-enhancing role of pQ. Instead, multiple SAR analyses support an indirect mechanism in which the N-terminal pQ lactam constrains ligand conformation and reweights a shared receptor-contact network. Extension to human NmU receptors further explain how the direction of such distal modulation is determined: the loose and permissive hNmU-R1 pocket better accommodates the constrained pQ ligand, whereas the compact hNmU-R2 pocket penalizes it. Thus, pQ converts a minimal chemical difference into subtype-specific GPCR signaling through receptor pocket-dependent conformational steering.

### Binding pocket permissiveness converts pQ constraint into subtype-selective signaling

Docking and mutagenesis analyses demonstrate that N-terminal pQ confers receptor subtype-selectivity via an indirect, conformation-based mechanism. Although MMG2-pDPb and MMG2-DPb engage largely overlapping receptor-contact residues, pQ reshapes the ligand’s bound configuration **(Fig. 2 C, D, Fig. 3 C, D, Fig. S13 C, and Fig. S18 A, B),** thereby altering the topology or strength of key interactions **(Fig. 2 E-G, Fig. 3 E-G and Fig. 4)**. Notably, the overall binding mode, in which C-terminus inserts into the orthosteric pocket while N-terminus remains solvent-exposed, closely parallels that observed in cryo-EM structures of human NmU-8 bound to its receptors (17, 18) **(Fig. S33)**, supporting a conserved directional binding architecture across phylogenetically distant neuropeptide-GPCR systems.

Classical “lock-key” model (49) is insufficient to capture the dynamic complexity of neuropeptide-GPCR interactions, as it neglects the intrinsic conformational flexibility of both ligands and receptors (47) **(Fig. 6 I)**. This limitation motivated the development of “induced fit” paradigm, which emphasizes receptor rearrangement following ligand engagement (44–47). However, given the diversity of neuropeptides and GPCR architectures, it remains unclear whether induced-fit applies universally. Importantly, “induced fit” models generally presume that optimization arises from direct ligand-receptor contacts (45, 63) and from a relatively extensive receptor binding pocket. Consequently, they do not readily explain how ligand features that do not directly contact the receptor can modulate signaling, nor why outcomes diverge across closely related receptor subtypes **(Fig.6 I)**.

Our findings address these issues by demonstrating that an N-terminal pQ moiety acts as a modulator to reshape the ligand core conformation, effectively reducing ligand flexibility **(Fig. 6 I)**. Importantly, the functional outcome of this constraint depends on receptor pocket permissiveness, where static pocket analysis and MD simulations further reveal that the underlying logic is governed by receptor pocket properties: the looser pocket of hNmU-R1 can accommodate the pQ-defined interaction pattern more productively than the non-pQ ligand, resulting in enhanced activation, resembling the canonical induced-fit response **(Fig. 4 and Fig. 6 I)**. Given the well-documented role of pQ in enhancing receptor activation across neuropeptide systems (23–26, 28), our results suggest that conformationally constrained ligands may be particularly effective in receptors that undergo induced-fit adaptations. In contrast, the more compact hNmU-R2 pocket cannot readily accommodate the pQ-imposed pattern relative to the non-pQ ligand, leading to conformational mismatch and suppressed activation **(Fig. 4 and Fig. 6 I)**. This represents an intermediate mode of recognition between static “lock-key” complementarity and the adaptive “induced-fit” accommodation, in which a ligand-borne conformational constraint is interpreted differently by receptor pockets of distinct permissiveness. This mode has not been formally described in neuropeptide-GPCR signaling and further implies that the receptors with compact pockets favor conformationally flexible ligands. Notably, these mechanisms likely explain the bidirectional regulation observed in the *Aplysia* PRXamide system.

We therefore define an indirect “PTM distal steering” mechanism, in which a non-contacting N-terminal PTM functions as a molecular tuner that reshapes the conserved ligand core to differentially engage receptor pockets of distinct architecture. Through this mechanism, a single peptide scaffold can generate opposed functional outputs across receptor subtypes via a minimal chemical change. In this framework, the *Aplysia* model reveals how pQ remotely reweights interaction networks at a molecular level **(Fig. 4)**, whereas the human NmU system further demonstrates how receptor pocket properties determine whether the pQ-constrained ligand is favored or penalized **(Fig. 6)**. This principle provides a mechanistic foundation for conformation-guided design of stable, subtype-selective peptide ligands and suggests that analogous distal conformational control may be broadly exploitable in small-molecule and macrocyclic ligand design to encode receptor selectivity without requiring direct contact modifications.

### Broader implications for PTM-mediated conformational control

Our findings position N-terminal pyroglutamylation within a broader class of PTMs that regulate GPCR signaling indirectly through conformational tuning. C-terminal amidation is well established to enhance receptor affinity through direct ligand-receptor interaction (16–19, 64). However, our prior work in *Aplysia* leucokinin system showed that the C-terminal amide does not directly penetrate the receptor pocket (65) (facing the solvent, as also observed in TRH (66)), yet its removal still abolishes receptor activation. This behavior parallels the conformational role delineated for pQ here, although precise mechanisms remained unsolved in that study. A related example is provided by *Aplysia* ATRP, in which L-to D-amino acid isomerization produces diastereomeric peptides with opposing effects on two receptor subtypes (67, 68), a phenomenon that may likewise arise from modification-induced conformational biasing. Similarly, N-terminal glycosylation in NPY and glucagon families may modulate GPCR signaling by altering peptide secondary structure or steric bulk (69). Disulfide bonds in oxytocin and vasopressin similarly confer ligand selectivity and stabilize productive ring conformations (3, 64, 70) without necessarily forming new receptor contacts (64), instead positioning internal residues for optimal interactions within a spatially constrained binding pocket.

Despite their chemical diversity, these PTMs converge on a shared functional logic: they impose topological constraints that shape ligand conformational landscapes and thereby tune interaction networks within the receptor pocket, rather than through direct contact alone.

### Conservation of pQ-dependent subtype-specific regulation and translational implications

The conservation of pQ-dependent modulation from *Aplysia* PRXamide to mammalian NmU signaling highlights its broader functional relevance. Although pQ-containing NmU peptides are rare, canine pQ-NmU-7 (pQFLFRPRNa) demonstrates that this modification can bias subtype preference in human NmU-Rs, consistent with the high sequence similarity between canine and human NmU receptors **(Fig. S34)**. The retained sensitivity of hNmU-R1 to pQ-mediated potentiation **(Fig. 6 A)** further supports the transferability of this chemical logic across species.

From a translational perspective, NmU receptors remain promising yet challenging therapeutic targets. hNmU-R2 has been prioritized for anti-obesity applications due to its metabolic efficacy with fewer side effects (71–74), whereas hNmU-R1 remains less explored but has been implicated in glucose-lowering effects (74, 75), immune regulation (76), and even potential application in treating bladder dysfunctions, including stress urinary incontinence or detrusor underactivity (77–80). However, clinical development of NmU-based ligands has been limited by peptide stability.

Although NmU peptides exhibit robust activity in receptor assays, their in vivo efficacy is constrained by rapid degradation (81–83). In this context, pQ provides a dual advantage: its known ability to enhance peptide stability (22, 24, 25) can be complemented by our finding that pQ can also tune receptor engagement through distal conformational control. Compared to synthetic N-terminal modifications or linker-based stabilization strategies (73, 84), pQ offers a chemically minimal, evolutionarily validated solution for improving both pharmacokinetic and pharmacodynamic properties.

Overall, our work defines an indirect “PTM distal steering” mechanism in which a non-contacting N-terminal pQ modulates subtype-specific GPCR activation by remodeling ligand conformation ensembles rather than by forming direct receptor contacts **(Fig. 4)**. The functional direction of this steering is dictated by receptor pocket mechanics: compact pockets favor more flexible ligands, whereas loose pockets better accommodate rigid pQ-constrained ligands **(Fig. 6 I)**. The extension of this molecular logic from *Aplysia* PRXamide to mammalian NmU signaling suggests that pQ can act as an evolutionarily conserved conformational switch capable of fine-tuning neuropeptide-GPCR communication. Given the widespread occurrence of N-terminal and cyclization PTMs, this mechanism may be relevant to other neuropeptide families and suggests that ligand-receptor interactions can operate through multiple regulatory modes beyond the canonical “induced-fit” paradigm. By translating this principle into a design framework, our study links fundamental mechanistic insight to new opportunities for the rational optimization of neuropeptide therapeutics, and may further inform the design of small-molecule and macrocyclic scaffolds.

## Materials and Methods

### Subjects and reagents

Experiments were performed on *Aplysia californica* (100-350 g) obtained from Marinus, California, USA. *Aplysia* are hermaphroditic (i.e., each animal has reproductive organs normally associated with both male and female sexes). Animals were maintained in circulating artificial seawater at 14°C-16°C and the animal room was equipped with a 24 h light cycle with the light period from 7:00 a.m. to 7:00 p.m. Before dissection, animals were anesthetized by injection of isotonic 333 mm MgCl_2_ (about 50% of body weight) into the body cavity. All reagents were purchased from Sigma-Aldrich unless otherwise stated. Peptides were synthesized by Sangon Biotech Pharmaceutical Co., Ltd, Guoping Pharmaceutical Co., Ltd., or Synpeptide Co., Ltd and the purity of the peptides was >95%. Peptides were aliquoted into 50 nmol per micro-centrifuge tube and stored at -20 °C until use.

### Peptide measurements by LC-MS/MS

To structurally characterize PRXamide peptides derived from the actual MMG2 precursor gene in the *Aplysia* CNS, peptide extracts from all central ganglia (abdominal, buccal, cerebral, pleural and pedal) of the CNS were sequenced *de novo* by LC-MS/MS (85). Isolated and desheathed ganglia were homogenized in acidified methanol (80:18:2 MeOH: water: acetic acid) using Precellys bead beater set to the “Soft” method (3 × 15 s, 5800 rpm protocol). Homogenates were collected and centrifuged at 14,000 x g for 15 min to precipitate tissue debris. Resulting supernatants were transferred into another set of vials, dried using a speed-vac. Dry crude extracts were reconstituted in 0.1% formic acid, mixed thoroughly, centrifuged, and the resulting supernatant was collected for further processing; the pellet was discarded.

*Aplysia* PRXamide peptides extracts were characterized using the Bruker nanoElute LC hyphenated to Bruker timsTOF Pro via CaptiveSpray source with external column oven (Bruker Daltonics, Billerica, MA). Samples were first injected onto Acclaim™ PepMap™ 100 C18 pre-column trap for additional desalting and preconcentration, followed by separation on a Bruker PepSep column (75 μm ID, 1.9 μm particle size, 250 mm length) at a flow rate of 400 nL/min and a column temperature of 35°C. The mobile phase consisted of solvent A (0.1% formic acid in water) and solvent B (0.1% formic acid in acetonitrile). A gradient of solvent B was applied, starting from 2% and increasing to 10% in 5 min and then to 50% over next 80 min, followed by a rapid ramp to 95% for wash, and finally equilibration with starting conditions. The positive ion mass spectra were acquired in Parallel Accumulation Serial Fragmentation (PASEF) mode with a cycle time of 1.1 s and 1/k0 range 0.6-1.6 V s/cm^2^.

### Identification of PRXamide receptor sequences in *Aplysia*

We primarily conducted sequence search and BLAST using the NCBI (https://www.ncbi.nlm.nih.gov/) and AplysiaTools (http://aplysiatools.org) databases. These latter databases (Dr Thomas Abrams, University of Maryland) include data for *Aplysia* transcriptome and genome (86). The ORFs of the putative receptor full-length cDNA sequence were obtained using ORF Finder (https://www.ncbi.nlm.nih.gov/orffinder/). BioEdit software (https://bioedit.software.informer.com/7.2/) was selected to compare the PRXa/NmU peptide and receptor sequences with those of other species. For the putative PRXamide receptors, TM domains and conserved sites were predicted using TMHMM Server, v.2.0 (87) (http://www.cbs.dtu.dk/services/TMHMM/) and NCBI Conserved Domain Database (88) (http://www.ncbi.nlm.nih. gov/Structure/cdd/cdd.shtml).

### Phylogenetic analysis and molecular evolution

Multiple sequence alignments of receptor sequences were performed by ClustalW and manually corrected. Phylogenetic trees were constructed by maximum likelihood (ML) methods of MEGA X and Phylosuite software. For Fig. S4, the LG+G+F model and 1000 times bootstrap were set. For Fig. S9, the LG+G model and 1000 times bootstrap were set. TBtools (89) and website Chiplot (90) (https://www.chiplot.online/tvbot.html) were used to combine and visualize the phylogenetic trees.

We used MUSCLE (codon) in MEGA X to compare the CDS sequences of PRXa/NmU receptors, and the aligned CDS sequences were converted into pml format for the construction of ML phylogenetic tree by Phylosuite (91) software. The ratio of non-synonymous/synonymous (ω) in the branch and specific positively selected sites was inferred using the Branch Model and Branch-Site Model in EasycodeML_v1.0 software (92), while Likelihood Ratio Test (LTR) was used to determine site differences (93).

### Generation of receptor plasmids

*Aplysia* PRXamide receptors were derived by RNA extraction and cDNA cloning. For RNA extraction, animals were anesthetized by injection of 333 mM MgCl_2_ (30-50% of body weight). Cerebral, pleural-pedal, buccal, and abdominal ganglia were then dissected and maintained in artificial seawater (460 mM NaCl, 10 mM KCl, 55 mM MgCl_2_, 11 mM CaCl_2_, 10 mM HEPES, pH 7.6) within Sylgard-lined dishes (Dow Corning). For RNA isolation, ganglia were transferred to 200 μL Trizol reagent (Sigma, T9424) and flash-frozen at -80°C.

Frozen samples were thawed and homogenized using a plastic pestle. Trizol volume was adjusted to 1 mL, followed by 10 min incubation at room temperature. Subsequent chloroform extraction (200 μL) involved vigorous vortexing and 15 min incubation on ice. Phase separation was achieved by centrifugation (12,000×g, 4°C, 15 min). The aqueous phase was combined with an equal volume of isopropanol, mixed by inversion, and precipitated at -20°C for 2 h. RNA pellets were obtained by centrifugation (12,000×g, 4°C, 15 min), washed twice with 75% ethanol, and air-dried for 5-10 min. Purified RNA was resuspended in 30 μL nuclease-free water, with concentration quantified via ND-1000 spectrophotometer (Thermo Fisher Scientific).

First-strand cDNA synthesis was performed using 1 μg total RNA with PrimeScript RT Master Mix (Takara, RR036A) under manufacturer-specified conditions. Synthesized cDNA was stored at -20°C until subsequent PCR.

The synthesized cDNA above was used as a polymerase chain reaction (PCR) template. Specific primers were designed based on protein-coding sequences for the putative ApPRXa-Rs. The PCR reaction was performed with 98 °C/2 min predenaturing, 98 °C/10 s denaturing, ∼64 °C/15 s annealing, 72 °C/30 s extension, and 72 °C/5 min re-extension for 35 cycles. The PCR products were subcloned into vector pcDNA3.1(+) or pcDNA3.1(+)-FLAG (for ELISA experiments only, see “Cell surface expression analysis by ELISA” section) and sequenced to ensure the sequences were correct.

Human Neuromedin U receptors (hNmU-R1 and hNmU-R2) were synthesized by Sangon Biotech Pharmaceutical Co., Ltd., based on sequences derived from the Genebank database (accession numbers: AF272362.1 for hNmU-R1 and AF272363.1 for hNmU-R2). The synthesized genes were subcloned into pcDNA3.1(+) and sequenced to ensure the sequences were correct.

### Cell culture and transfection

The CHO-K1 cells (Procell, CL-0062) were cultured in F-12K medium (Gibco, 21127-022) with 10% fetal bovine serum (FBS) (Genial, G11-70500) at 37 °C in a humidified incubator containing 5% CO_2_. Cell culture dishes (Corning, 430166) were used for culture until cells reached the logarithmic growth phase. Then, the medium was discarded, and cells were washed twice with PBS and digested with trypsin. Transfection experiments were performed when the cells were grown to 70−90% confluence and using jetPRIME (Polyplus Transfection, PT-114-15) transfection reagent, according to the manufacturer’s instructions. Specifically, in all experiments in this study, for each 35mm dish, 2 μg of the putative receptor plasmids [in pcDNA3.1(+) or pcDNA3.1(+)-FLAG] and 2 μg Gαq plasmids [in pcDNA3.1(+)] were mixed with 200 μL jetPRIME Buffer (Lot# 0000000396), followed by the addition of 6 μL jetPRIME. Gαq plasmids were added to ensure that the ligand receptor binding would generate an IP1 response (see “IP1 accumulation assay section”) no matter what Gα protein a putative receptor might associate because Gαq is a promiscuous protein that will bind to most GPCRs. The CHO cells with the reagents added above were mixed gently and incubated at room temperature for 15 min. The DNA/jetPRIME mixture dropwise was then added to the dish, and the cells were incubated at 37 °C in 5% CO_2_ overnight.

### IP1 accumulation assay

Inositol monophosphate (IP1) accumulation assay measures the concentration of IP1, which is hydrolyzed from the second messenger, inositol trisphosphate (IP3), generated by the Gαq pathway (94, 95) when a G-protein coupled receptor (GPCR) expressed in CHO-K1 cells is activated by an appropriate ligand. On the third day, transfected cells were trypsinized and reseeded in 384-well tissue culture-treated plates (Corning, 3570) at a density of 20,000 cells/well in F-12K and 10% FBS at 37 °C in 5% CO_2_ overnight. The next day, the medium in wells was discarded and exposed to potential peptide agonists in 14 μL stimulation buffer (Lot No. 621P1FDC) for 1h. Peptide agonists, ranging in concentrations from 10^−12^ to 10^−4^ M, were used to construct dose-response curves. The activation of the putative receptors was detected by monitoring intracellular IP1 concentration using an IP-One Gq kit (Cisbio, 62IPAPEB) in Tecan Spark following the manufacturer’s instructions with minor modifications (for using 0.5×reagents for the anti-IP1-cryptate and IP1-d2 reagents). Specifically, after adding the detection solution to each well, the sealed plate was incubated for a minimum of 1h at RT protected from light, and each plate was then read with Spark (TECAN, USA) with an HTRF compatible setup: excitation at 320 nm and emitted light at 620 and 665 nm.

### Molecular modeling and docking

Since pyroglutamic acid (pQ) is a non-canonical amino acid, MMG2-pDPb/DPb and other ligands were constructed using ChemDraw (96) and exported in sdf format for subsequent molecular docking analysis. No manual energy minimization was applied, as all docking protocols used in this study perform internal conformational sampling and scoring. Three-dimensional models of all receptors were generated independently using both the website Swiss model (97) (https://swissmodel.expasy.org/) and the Robetta server (98) (http://robetta.bakerlab.org/), the resulting models were compared for overall structural consistency. Model qualities were assessed using the SAVES v6.1 server (https://saves.mbi.ucla.edu/), and stereochemical properties were evaluated by Ramachandran plot analysis with MolProbity (99) (https://molprobity.biochem.duke.edu/).

Peptide-protein docking was performed primarily using HPEPDOCK 2.0 (56) (http://huanglab.phys.hust.edu.cn/hpepdock/) and diffdock (57) (https://huggingface.co/spaces/simonduerr/diffdock). HPEPDOCK supports the input of peptide sequences containing non-standard amino acids; accordingly, ligand sequences were submitted directly with the pre-modeled receptor structures. In contrast, diffdock requires explicit structural input, including ligand sdf files and receptor pdb files. For each docking output, blind docking was performed, and the top 10 ranked models with plausible binding pocket engagement were selected for interaction analysis. Recurrent interacting residues and representative docking poses were visualized and analyzed using PyMOL and ChimeraX (100, 101). Additional detailed docking workflow and complementary peptide-protein docking are provided in the **Supplementary Results and Discussion** section.

### Molecular dynamics simulations

The CHARMM-GUI platform (102) was employed to integrate the structures into a hexagon pure 1-palmitoyl-2-oleoyl-sn-glycerol-3-phosphocholine (POPC) lipid bilayer, solvated with a 22.5 Å water layer and 150 mM Na^+^/Cl^-^. Given that MMG2-pDPb contains an N-terminal pQ residue that is not included in the PTM library of CHARMM-GUI, we excluded the predefined amino acid annotation of the ligand peptide and performed atomistic modeling by MOE (103) to retain and simulate the pQ moiety. All systems were described using the CHARMM36m (104) force field, and all MD simulations were carried out using NAMD 3 (105). Each system was first subjected to 5000 steps of energy minimization, followed by the standard six-step equilibration protocol as recommended by CHARMM-GUI to gradually relax the membrane, solvent and protein components. Production simulations were then performed for 300 ns under constant temperature (303.15 K) and pressure (1 bar**) (Fig. 5 E)**. The stability of the simulated systems was evaluated by calculating the root-mean-square deviation (RMSD), root-mean-square fluctuation (RMSF), free energy surface (FES), and radius of gyration (Rg) based on all heavy atoms of the receptor-peptide complexes. Trajectory analysis and all calculations were conducted using VMD (106) and MDAnalysis (107).

### Binding-pocket volume analysis

The orthosteric binding pockets of human NmU receptors were defined and quantified using CASTp with a probe radius of 1.4 Å (62). Ligand-bound active-state structures of hNmU-R1 and hNmU-R2 were used for pocket analysis (17, 18). Before CASTp calculation, non-receptor components, including NmU-8, G proteins, lipids, and solvent molecules, were removed, while the ligand-bound receptor conformations were retained. Molecular surface volume (MS volume) was used to compare pocket size between receptor subtypes. Structural visualization of receptor pockets and CASTp-defined pocket surfaces was performed in UCSF Chimera (108).

### Mutagenesis of the ApPRXa-Rs

In a first set of experiments, mutagenesis of the receptors was performed without a FLAG tag. Specifically, construction of the mutants was performed employing the full-length ApPRXa-Rs cDNA cloned into the pcDNA3.1(+) plasmid. Site-directed mutagenesis was performed using the site-directed mutagenesis kit (Sangon Biotech), following the manufacturer’s instructions. Briefly, forward and reverse primers containing the expected mutation were mixed with kit components, and 10 ng of pcDNA3.1(+)-receptors was used as the mutation template. After 14 to 18 rounds of PCR amplification, 1 μl of DpnI was added and incubated at 37 °C for 1 h to digest the template. The primers used to obtain the mutants were designed based on the cDNA sequences as listed in Supplementary Table 2. Mutants were confirmed via DNA sequencing. IP1 accumulation assay with ApPRXa-Rs and mutants without FLAG tags was performed using procedures described in “IP1 accumulation assay” section.

### Cell surface expression analysis by ELISA

Cell surface expression determination of ApPRXa-Rs and mutant receptors, all with FLAG tags, was performed using a procedure modified from previous work (23, 65, 109–111). Specifically, CHO-K1 cells were maintained in F-12K medium with 10% FBS at 37 °C in 5% CO_2_. When the cells were grown to 70 to 90% confluence, transfection experiments were performed. Briefly, in each well, 2 μg of the receptor plasmids [in pcDNA3.1(+)-FLAG] were mixed with 200 μl of jetPRIME buffer, followed by the addition of 6 μl of jetPRIME and incubated at room temperature for 15 min. The DNA/jetPRIME mixture dropwise was then added to the dish. The cells were incubated at 37 °C in 5% CO_2_ overnight. After 24 h, transfected cells were plated in 96-well white clear-bottom cell culture plates (Corning, 3610) at a density of 20,000 cells in 100 μl per well and incubated overnight. The following day, culture media was aspirated and cells were washed twice with 200 μlof1×PBS (Procell, PB180327). Then, 100 μl of 1× PBS containing 5% (w/v) bovine serum albumin was added to each well and incubated at room temperature. After 30 min, 100 μl of 1:10,000 anti-FLAG M2-HRP conjugate (Sigma-Aldrich, Cat A8592) was added to each well and incubated for 30 min at 37 °C. Cells were washed twice with 200 μl of 1× PBS and then incubated with 200 μl of TMB Chromogen Solution (Sangon Biotech, E661007) for 30 min at 37 °C in the dark. Finally, 50 μl of ELISA Stopping Solution (Sangon Biotech, E661006) was added to each well to stop the reaction. The absorbance at 450 nm was measured using a Tecan Spark microplate reader. In each experiment, expression of the mutant was assessed and compared with that measured for the corresponding wildtype receptor in the same experiment.

### Statistical analysis

Dose-response curves and bar graphs for experimental data were plotted using Prism software (GraphPad, https://www. graphpad.com/). Data are presented as mean ± SEM of at least three independent experiments. Statistical tests were performed using Prism software. They included Student’s t-test, and one-way or two-way ANOVA, as appropriate. Statistical significance was set at P < 0.05.

## Acknowledgements

This work was supported by the National Natural Science Foundation of China (grants 32471073, 32171011, 31861143036, 31671097 and 31371104 to J. J., grant 32100816 to G.Z.), the Department of Chemistry (UIUC) Discovery Fund to J. V. S., and the Natural Science Foundation of Henan Province (grant 262300421110 to G. Z.). We thank the Collaborative Innovation Center of Advanced Microstructures, Nanjing University, for providing the computational resources and technical support required for the molecular dynamics simulations.

## Additional information

This article contains supporting information, which includes Supporting Results and Discussion, Table S1-9, and Figs. S1-34.

## Author contributions

J.-h. C., W.-j. L., and J. J. writing–original draft; J.-h. C., W.-j. L., G. Z., S.-q. W., C.-p. L., J. V. S., and E. V. R. validation; J.-h. C., W.-j. L., S.-q. W., W.-j. L., C.-p. L., F. L., X.-y. D., H.-y. W., J.-p. X., P. F., Y.-l. Z., Q.-c. J., C.-y. L., R.-t. M., Y.-c.-f. Z., G. Z., J. V. S., E. V. R. and J. J. investigation; J.-h. C., W.-j. L., S.-q. W., C.-p. L., J. V. S., E. V. R. data curation; J.-h. C., W.-j. L., and J. J. conceptualization; J.-h. C., W.-j. L., G. Z., J. V. S., E. V. R., and J. J. writing– review and editing; G. Z., J. V. S., and J. J. funding acquisition; J. J. supervision; J. J. project administration; J. J. methodology.

## References

1. Craig, A. G., Bandyopadhyay, P., and Olivera, B. M. (1999) Post-translationally modified neuropeptides from *Conus venoms*. European Journal of Biochemistry 264, 271–275

2. Bai, L., Livnat, I., Romanova, E. V., Alexeeva, V., Yau, P. M., Vilim, F. S. et al. (2013) Characterization of GdFFD, a D-amino acid-containing neuropeptide that functions as an extrinsic modulator of the *Aplysia* feeding circuit. Journal of Biological Chemistry 288, 32837–32851

3. Xu, J.-P., Ding, X.-Y., Guo, S.-Q., Wang, H.-Y., Liu, W.-J., Jiang, H.-M. et al. (2023) Characterization of an *Aplysia* vasotocin signaling system and actions of posttranslational modifications and individual residues of the ligand on receptor activity. Frontiers in Pharmacology 14, 1132066

4. Chang, J.-H., Wu, S.-Q., Ding, X.-Y., Pan, X., Wang, H.-Y., He, Y.-X. et al. (2026) Decoding of amidated aromatic C-terminus and sulfation by cholecystokinin receptors reveals conserved and divergent evolutionary mechanisms. Journal of Biological Chemistry 10.1016/j.jbc.2026.113327

5. Chen, R., Hui, L., Cape, S. S., Wang, J., and Li, L. (2010) Comparative neuropeptidomic analysis of food intake via a multifaceted mass spectrometric approach. ACS chemical neuroscience 1, 204–214

6. Nusbaum, M. P., and Blitz, D. M. (2012) Neuropeptide modulation of microcircuits. Current Opinion in Neurobiology 22, 592–601

7. Taghert, P. H., and Nitabach, M. N. (2012) Peptide neuromodulation in invertebrate model systems. Neuron 76, 82–97

8. Martinez, V. G., and O’Driscoll, L. (2015) Neuromedin U: a multifunctional neuropeptide with pleiotropic roles. Clinical Chemistry 61, 471–482

9. Cropper, E. C., Jing, J., Vilim, F. S., Barry, M. A., and Weiss, K. R. (2018) Multifaceted expression of peptidergic modulation in the feeding system of *Aplysia*. ACS Chemical Neuroscience 9, 1917–1927

10. Abid, M. S. R., Mousavi, S., and Checco, J. W. (2021) Identifying receptors for neuropeptides and peptide hormones: challenges and recent progress. ACS Chemical Biology 16, 251–263

11. Zhang, G., Guo, S., Wang, H., Li, Y., Jiang, H., and Yu, K. (2022) Functional studies on neuropeptides and receptors in model animal *Aplysia* (in Chinese). Sci. Sin Vitae 52, 387–402

12. Li, K., Shi, W., Tan, Y., Ding, Y., Armstrong, A. G., Vlasov, Y., and Sweedler, J. V. (2025) Neuropeptides in the extracellular space of the mouse cortex measured in vivo by nanodialysis probe coupled with LC-MS. Angewandte Chemie 137, e202509490

13. Lentschat, H., Liessmann, F., Tydings, C., Schermeng, T., Stichel, J., Urban, N. et al. (2025) Hederagenin is a highly selective antagonist of the neuropeptide FF receptor 1 that reveals mechanisms for subtype selectivity. Angewandte Chemie 137, e202417786

14. Escher, E., Couture, R., Poulos, C., Pinas, N., Mizrahi, J., Theodoropoulos, D., and Regoli, D. (1982) Structure-activity studies on the C-terminal amide of substance P. Journal of Medicinal Chemistry 25, 1317–1321

15. Liu, Q., Yang, D., Zhuang, Y., Croll, T. I., Cai, X., Dai, A. et al. (2021) Ligand recognition and G-protein coupling selectivity of cholecystokinin A receptor. Nature chemical biology 17, 1238–1244

16. Park, C., Kim, J., Ko, S.-B., Choi, Y. K., Jeong, H., Woo, H. et al. (2022) Structural basis of neuropeptide Y signaling through Y1 receptor. Nature Communications 13, 853

17. You, C., Zhang, Y., Xu, P., Huang, S., Yin, W., Eric Xu, H., and Jiang, Y. (2022) Structural insights into the peptide selectivity and activation of human neuromedin U receptors. Nature Communications 13, 2045

18. Zhao, W., Zhang, W., Wang, M., Lu, M., Chen, S., Tang, T. et al. (2022) Ligand recognition and activation of neuromedin U receptor 2. Nature Communications 13, 7955

19. Iwama, A., Kise, R., Akasaka, H., Sano, F. K., Oshima, H. S., Inoue, A. et al. (2024) Structure and dynamics of the pyroglutamylated RF-amide peptide QRFP receptor GPR103. Nature Communications 15, 4769

20. Cynis, H., Schilling, S., Bodnár, M., Hoffmann, T., Heiser, U., Saido, T. C., and Demuth, H.-U. (2006) Inhibition of glutaminyl cyclase alters pyroglutamate formation in mammalian cells. Biochimica et Biophysica Acta (BBA)-Proteins and Proteomics 1764, 1618–1625

21. Kumar, A., and Bachhawat, A. K. (2012) Pyroglutamic acid: throwing light on a lightly studied metabolite. Current Science 288–297

22. Wiggenhorn, A. L., Abuzaid, H. Z., Coassolo, L., Li, V. L., Tanzo, J. T., Wei, W. et al. (2023) A class of secreted mammalian peptides with potential to expand cell-cell communication. Nature Communications 14, 8125

23. Mao, R.-t., Guo, S.-q., Zhang, G., Li, Y.-d., Xu, J.-p., Wang, H.-y., et al. (2024) Two C-terminal isoforms of *Aplysia* tachykinin–related peptide receptors exhibit phosphorylation-dependent and phosphorylation-independent desensitization mechanisms. Journal of Biological Chemistry 300, 107556

24. Abraham, G. N., and Podell, D. N. (1981) Pyroglutamic acid: non-metabolic formation, function in proteins and peptides, and characteristics of the enzymes effecting its removal. Molecular and Cellular Biochemistry 38, 181–190

25. Morty, R. E., Bulau, P., Pellé, R., Wilk, S., and Abe, K. (2006) Pyroglutamyl peptidase type I from Trypanosoma brucei: a new virulence factor from African trypanosomes that de-blocks regulatory peptides in the plasma of infected hosts. Biochemical Journal 394, 635–645

26. Hashimoto, T., Kurosawa, K., and Sakura, N. (1995) Structure-activity relationships of neuromedin U. II. Highly potent analogs substituted or modified at the N-terminus of neuromedin U-8. Chemical and Pharmaceutical Bulletin 43, 1154–1157

27. Chung, J. S., and Webster, S. G. (1996) Does the N-terminal pyroglutamate residue have any physiological significance for crab hyperglycemic neuropeptides? European Journal of Biochemistry 240, 358–364

28. Howl, J., Yarwood, N., Stock, D., and Wheatley, M. (1996) Probing the V1a vasopressin receptor binding site with pyroglutamate-substituted linear antagonists. Neuropeptides 30, 73–79

29. Kreienkamp, H.-J., Larusson, H. J., Witte, I., Roeder, T., Birgul, N., Hönck, H.-H. et al. (2002) Functional annotation of two orphan G-protein-coupled receptors, Drostar1 and-2, from *Drosophila melanogaster* and their ligands by reverse pharmacology. Journal of Biological Chemistry 277, 39937–39943

30. Jackson, G. E., and Gäde, G. (2021) Insights into the activation of a crustacean G protein-coupled receptor: evaluation of the red pigment-concentrating hormone receptor of the water flea *Daphnia pulex* (Dappu-RPCH R) Biomolecules. 11, 710

31. Yeung, H. Y., Ramiro, I. B. L., Andersen, D. B., Koch, T. L., Hamilton, A., Bjørn-Yoshimoto, W. E. et al. (2024) Fish-hunting cone snail disrupts prey’s glucose homeostasis with weaponized mimetics of somatostatin and insulin. Nature Communications 15, 6408

32. Gilon, C., Halle, D., Chorev, M., Selincer, Z., and Byk, G. (1991) Backbone cyclization: a new method for conferring conformational constraint on peptides. Biopolymers: Original Research on Biomolecules 31, 745–750

33. Nath, S., Buell, A. K., and Barz, B. (2023) Pyroglutamate-modified amyloid β (3–42) monomer has more β-sheet content than the amyloid β (1–42) monomer. Physical Chemistry Chemical Physics 25, 16483–16491

34. Marco, H. G., Verlinden, H., Vanden Broeck, J., and Gäde, G. (2017) Characterisation and pharmacological analysis of a crustacean G protein-coupled receptor: the red pigment-concentrating hormone receptor of *Daphnia pulex*. Scientific reports 7, 6851

35. Minamino, N., Kangawa, K., and Matsuo, H. (1985) Neuromedin U-8 and U-25: novel uterus stimulating and hypertensive peptides identified in porcine spinal cord. Biochemical and biophysical research communications 130, 1078–1085

36. Tublitz, N. J., and Truman, J. W. (1985) Insect cardioactive peptides: I. Distribution and molecular characteristics of two cardioacceleratory peptides in the tobacco hawkmoth, *Manduca sexta*. Journal of experimental biology 114, 365–379

37. Holman, G., Cook, B., and Nachman, R. (1986) Primary structure and synthesis of a blocked myotropic neuropeptide isolated from the cockroach, *Leucophaea maderae*. *Comparative biochemistry and physiology. C*, Comparative pharmacology and toxicology 85, 219–224

38. Park, Y., Kim, Y.-J., and Adams, M. E. (2002) Identification of G protein-coupled receptors for *Drosophila* PRXamide peptides, CCAP, corazonin, and AKH supports a theory of ligand-receptor coevolution. Proceedings of the National Academy of Sciences 99, 11423–11428

39. Saideman, S. R., Ma, M., Kutz-Naber, K. K., Cook, A., Torfs, P., Schoofs, L. et al. (2007) Modulation of rhythmic motor activity by pyrokinin peptides. Journal of neurophysiology 97, 579–595

40. Proekt, A., Vilim, F. S., Alexeeva, V., Brezina, V., Friedman, A., Jing, J. et al. (2005) Identification of a new neuropeptide precursor reveals a novel source of extrinsic modulation in the feeding system of *Aplysia*. Journal of Neuroscience 25, 9637–9648

41. Beuming, T., and Sherman, W. (2012) Current assessment of docking into GPCR crystal structures and homology models: successes, challenges, and guidelines. Journal of Chemical Information and Modeling 52, 3263–3277

42. Weng, G., Gao, J., Wang, Z., Wang, E., Hu, X., Yao, X. et al. (2020) Comprehensive evaluation of fourteen docking programs on protein–peptide complexes. Journal of Chemical Theory and Computation 16, 3959–3969

43. O’Harte, F., Bockman, C. S., Abel, P. W., and Conlon, J. M. (1991) Isolation, structural characterization and pharmacological activity of dog neuromedin U. Peptides 12, 11–15

44. Koshland Jr, D. E. (1994) The key–lock theory and the induced fit theory. Angewandte Chemie International Edition in English 33, 2375–2378

45. Bosshard, H. R. (2001) Molecular recognition by induced fit: how fit is the concept? News in Physiological Sciences 16, 171–173

46. Tobi, D., and Bahar, I. (2005) Structural changes involved in protein binding correlate with intrinsic motions of proteins in the unbound state. Proceedings of the National Academy of Sciences 102, 18908–18913

47. A Sotriffer, C. (2011) Accounting for induced-fit effects in docking: what is possible and what is not? Current Topics in Medicinal Chemistry 11, 179–191

48. Krumm, B. E., and Grisshammer, R. (2015) Peptide ligand recognition by G protein-coupled receptors. Frontiers in Pharmacology 6, 48

49. Fischer, E. (1894) Einfluss der Configuration auf die Wirkung der Enzyme. Berichte der deutschen chemischen Gesellschaft 27, 2985–2993

50. Zestos, A. G., and Kennedy, R. T. (2017) Microdialysis coupled with LC-MS/MS for in vivo neurochemical monitoring. The AAPS journal 19, 1284–1293

51. Neumann, E. K., Do, T. D., Comi, T. J., and Sweedler, J. V. (2019) Exploring the fundamental structures of life: non-targeted, chemical analysis of single cells and subcellular structures. Angewandte Chemie International Edition 58, 9348–9364

52. Coleman, J. L., Ngo, T., and Smith, N. J. (2017) The G protein-coupled receptor N-terminus and receptor signalling: N-tering a new era. Cellular signalling 33, 1–9

53. Rajagopal, S., and Shenoy, S. K. (2018) GPCR desensitization: Acute and prolonged phases. Cellular signalling 41, 9–16

54. Smith, J. S., Lefkowitz, R. J., and Rajagopal, S. (2018) Biased signalling: from simple switches to allosteric microprocessors. Nature reviews Drug discovery 17, 243–260

55. Patwardhan, A., Cheng, N., and Trejo, J. (2021) Post-translational modifications of g protein– coupled receptors control cellular signaling dynamics in space and time. Pharmacological Reviews 73, 120–151

56. Zhou, P., Jin, B., Li, H., and Huang, S.-Y. (2018) HPEPDOCK: a web server for blind peptide– protein docking based on a hierarchical algorithm. Nucleic Acids Research 46, W443–W450

57. Corso, G., Stärk, H., Jing, B., Barzilay, R., and Jaakkola, T. (2022) Diffdock: Diffusion steps, twists, and turns for molecular docking. arXiv preprint arXiv:2210.01776

58. Shonberg, J., Kling, R. C., Gmeiner, P., and Löber, S. (2015) GPCR crystal structures: medicinal chemistry in the pocket. Bioorganic & Medicinal Chemistry 23, 3880–3906

59. Black, S. D., and Mould, D. R. (1991) Development of hydrophobicity parameters to analyze proteins which bear post-or cotranslational modifications. Analytical biochemistry 193, 72–82

60. Ballesteros, J. A., Jensen, A. D., Liapakis, G., Rasmussen, S. G., Shi, L., Gether, U., and Javitch, J. A. (2001) Activation of the β2-adrenergic receptor involves disruption of an ionic lock between the cytoplasmic ends of transmembrane segments 3 and 6. Journal of Biological Chemistry 276, 29171–29177

61. Kawai, T., Katayama, Y., Guo, L., Liu, D., Suzuki, T., Hayakawa, K. et al. (2014) Identification of functionally important residues of the silkmoth pheromone biosynthesis-activating neuropeptide receptor, an insect ortholog of the vertebrate neuromedin U receptor. Journal of Biological Chemistry 289, 19150–19163

62. Ye, B., Tian, W., Wang, B., and Liang, J. (2024) CASTpFold: Computed atlas of surface topography of the universe of protein folds. Nucleic Acids Research 52, W194–W199

63. Di Cera, E. (2020) Mechanisms of ligand binding. Biophysics reviews 1,

64. Zhou, F., Ye, C., Ma, X., Yin, W., Croll, T. I., Zhou, Q. et al. (2021) Molecular basis of ligand recognition and activation of human V2 vasopressin receptor. Cell research 31, 929–931

65. Guo, S.-Q., Li, Y.-D., Chen, P., Zhang, G., Wang, H.-Y., Jiang, H.-M. et al. (2022) AI protein structure prediction-based modeling and mutagenesis of a protostome receptor and peptide ligands reveal key residues for their interaction. Journal of Biological Chemistry 298,

66. Yang, F., Zhang, H., Meng, X., Li, Y., Zhou, Y., Ling, S. et al. (2022) Structural insights into thyrotropin-releasing hormone receptor activation by an endogenous peptide agonist or its orally administered analogue. Cell Research 32, 858–861

67. Checco, J. W., Zhang, G., Yuan, W.-d., Le, Z.-w., Jing, J., and Sweedler, J. V. (2018) *Aplysia* allatotropin-related peptide and its newly identified D-amino acid–containing epimer both activate a receptor and a neuronal target. Journal of Biological Chemistry 293, 16862–16873

68. Yussif, B. M., Blasing, C. V., and Checco, J. W. (2023) Endogenous L-to D-amino acid residue isomerization modulates selectivity between distinct neuropeptide receptor family members. Proceedings of the National Academy of Sciences 120, e2217604120

69. Madsen, T. D., Hansen, L. H., Hintze, J., Ye, Z., Jebari, S., Andersen, D. B. et al. (2020) An atlas of O-linked glycosylation on peptide hormones reveals diverse biological roles. Nature Communications 11, 4033

70. Antobreh, G., Enyedy, I., and Ravna, A. W. (2018) Molecular modeling and docking studies of the oxytocin receptor. Future Medicinal Chemistry 10, 135–155

71. Kaisho, T., Nagai, H., Asakawa, T., Suzuki, N., Fujita, H., Matsumiya, K. et al. (2017) Effects of peripheral administration of a Neuromedin U receptor 2-selective agonist on food intake and body weight in obese mice. International Journal of Obesity 41, 1790–1797

72. Kanematsu-Yamaki, Y., Nishizawa, N., Kaisho, T., Nagai, H., Mochida, T., Asakawa, T. et al. (2017) Potent body weight-lowering effect of a neuromedin U receptor 2-selective PEGylated peptide. Journal of medicinal chemistry 60, 6089–6097

73. Nishizawa, N., Kanematsu-Yamaki, Y., Funata, M., Nagai, H., Shimizu, A., Fujita, H. et al. (2017) A potent neuromedin U receptor 2-selective alkylated peptide. Bioorganic & Medicinal Chemistry Letters 27, 4626–4629

74. Nagai, H., Kaisho, T., Yokoyama, K., Asakawa, T., Fujita, H., Matsumiya, K. et al. (2018) Differential effects of selective agonists of neuromedin U1 and U2 receptors in obese and diabetic mice. British Journal of Pharmacology 175, 359–373

75. Meier, J. J. (2012) GLP-1 receptor agonists for individualized treatment of type 2 diabetes mellitus. Nature Reviews Endocrinology 8, 728–742

76. Klose, C. S., Mahlakõiv, T., Moeller, J. B., Rankin, L. C., Flamar, A.-L., Kabata, H. et al. (2017) The neuropeptide neuromedin U stimulates innate lymphoid cells and type 2 inflammation. Nature 549, 282–286

77. Maggi, C. A., Patacchini, R., Giuliani, S., Turini, D., Barbanti, G., Rovero, P., and Meli, A. (1990) Motor response of the human isolated small intestine and urinary bladder to porcine neuromedin U-8. British journal of pharmacology 99, 186

78. Westfall, T. D., McCafferty, G. P., Pullen, M., Gruver, S., Sulpizio, A. C., Aiyar, V. N. et al. (2002) Characterization of neuromedin U effects in canine smooth muscle. The Journal of pharmacology and experimental therapeutics 301, 987–992

79. Luber, K. M. (2004) The definition, prevalence, and risk factors for stress urinary incontinence. Reviews in urology 6, S3

80. Osman, N. I., and Chapple, C. R. (2014) Contemporary concepts in the aetiopathogenesis of detrusor underactivity. Nature Reviews Urology 11, 639–648

81. Peier, A. M., Desai, K., Hubert, J., Du, X., Yang, L., Qian, Y. et al. (2011) Effects of peripherally administered neuromedin U on energy and glucose homeostasis. Endocrinology 152, 2644–2654

82. Dalbøge, L. S., Pedersen, S. L., van Witteloostuijn, S. B., Rasmussen, J. E., Rigbolt, K. T., Jensen, K. J., et al. (2015) Synthesis and evaluation of novel lipidated neuromedin U analogs with increased stability and effects on food intake. Journal of Peptide Science 21, 85–94

83. Masuda, Y., Kumano, S., Noguchi, J., Sakamoto, K., Inooka, H., and Ohtaki, T. (2017) PEGylated neuromedin U-8 shows long-lasting anorectic activity and anti-obesity effect in mice by peripheral administration. Peptides 94, 99–105

84. De Prins, A., Martin, C., Van Wanseele, Y., Skov, L. J., Tömböly, C., Tourwé, D., et al. (2018) Development of potent and proteolytically stable human neuromedin U receptor agonists. European journal of medicinal chemistry 144, 887–897

85. Tran, N. H., Qiao, R., Xin, L., Chen, X., Liu, C., Zhang, X. et al. (2019) Deep learning enables de novo peptide sequencing from data-independent-acquisition mass spectrometry. Nature methods 16, 63–66

86. Orvis, J., Albertin, C. B., Shrestha, P., Chen, S., Zheng, M., Rodriguez, C. J. et al. (2022) The evolution of synaptic and cognitive capacity: Insights from the nervous system transcriptome of *Aplysia*. Proceedings of the National Academy of Sciences 119, e2122301119

87. Krogh, A., Larsson, B., Von Heijne, G., and Sonnhammer, E. L. (2001) Predicting transmembrane protein topology with a hidden Markov model: application to complete genomes. Journal of Molecular Biology 305, 567–580

88. Marchler-Bauer, A., Derbyshire, M. K., Gonzales, N. R., Lu, S., Chitsaz, F., Geer, L. Y. et al. (2015) CDD: NCBI’s conserved domain database. Nucleic acids research 43, D222–D226

89. Chen, C., Chen, H., Zhang, Y., Thomas, H. R., Frank, M. H., He, Y., and Xia, R. (2020) TBtools: an integrative toolkit developed for interactive analyses of big biological data. Molecular plant 13, 1194–1202

90. Xie, J., Chen, Y., Cai, G., Cai, R., Hu, Z., and Wang, H. (2023) Tree Visualization By One Table (tvBOT): a web application for visualizing, modifying and annotating phylogenetic trees. Nucleic acids research 51, W587–W592

91. Zhang, D., Gao, F., Jakovlić, I., Zou, H., Zhang, J., Li, W. X., and Wang, G. T. (2020) PhyloSuite: An integrated and scalable desktop platform for streamlined molecular sequence data management and evolutionary phylogenetics studies. Molecular ecology resources 20, 348–355

92. Gao, F., Chen, C., Arab, D. A., Du, Z., He, Y., and Ho, S. Y. (2019) EasyCodeML: A visual tool for analysis of selection using CodeML. Ecology and evolution 9, 3891–3898

93. Ou, S., Chen, J., and Jiang, N. (2018) Assessing genome assembly quality using the LTR Assembly Index (LAI). Nucleic acids research 46, e126–e126

94. Bauknecht, P., and Jékely, G. (2015) Large-scale combinatorial deorphanization of *Platynereis* neuropeptide GPCRs. Cell Reports 12, 684–693

95. Sharma, S., and Checco, J. W. (2021) Evaluating functional ligand-GPCR interactions in cell-based assays In Methods in Cell Biology, Elsevier, 15–42

96. Mendelsohn, L. D. (2004) ChemDraw 8 ultra, windows and macintosh versions. Journal of Chemical Information and Computer Sciences 44, 2225–2226

97. Schwede, T., Kopp, J., Guex, N., and Peitsch, M. C. (2003) SWISS-MODEL: an automated protein homology-modeling server. Nucleic Acids Research 31, 3381–3385

98. Kim, D. E., Chivian, D., and Baker, D. (2004) Protein structure prediction and analysis using the Robetta server. Nucleic Acids Research 32, W526–W531

99. Chen, V. B., Arendall, W. B., Headd, J. J., Keedy, D. A., Immormino, R. M., Kapral, G. J. et al. (2010) MolProbity: all-atom structure validation for macromolecular crystallography. Biological Crystallography 66, 12–21

100. DeLano, W. L. (2002) Pymol: An open-source molecular graphics tool. CCP4 Newsl. Protein Crystallogr 40, 82–92

101. Pettersen, E. F., Goddard, T. D., Huang, C. C., Meng, E. C., Couch, G. S., Croll, T. I. et al. (2021) UCSF ChimeraX: Structure visualization for researchers, educators, and developers. Protein Science 30, 70–82

102. Jo, S., Kim, T., Iyer, V. G., and Im, W. (2008) CHARMM-GUI: a web-based graphical user interface for CHARMM. Journal of computational chemistry 29, 1859–1865

103. Vilar, S., Cozza, G., and Moro, S. (2008) Medicinal chemistry and the molecular operating environment (MOE): application of QSAR and molecular docking to drug discovery. Current Topics in Medicinal Chemistry 8, 1555–1572

104. Huang, J., Rauscher, S., Nawrocki, G., Ran, T., Feig, M., De Groot, B. L., et al. (2017) CHARMM36m: an improved force field for folded and intrinsically disordered proteins. Nature methods 14, 71–73

105. Phillips, J. C., Braun, R., Wang, W., Gumbart, J., Tajkhorshid, E., Villa, E. et al. (2005) Scalable molecular dynamics with NAMD. Journal of computational chemistry 26, 1781–1802

106. Humphrey, W., Dalke, A., and Schulten, K. (1996) VMD: visual molecular dynamics. Journal of molecular graphics 14, 33–38

107. Gowers, R. J., Linke, M., Barnoud, J., Reddy, T. J. E., Melo, M. N., Seyler, S. L., et al. (2019) MDAnalysis: a Python package for the rapid analysis of molecular dynamics simulations Los Alamos National Laboratory (LANL), Los Alamos, NM (United States),

108. Pettersen, E. F., Goddard, T. D., Huang, C. C., Couch, G. S., Greenblatt, D. M., Meng, E. C., and Ferrin, T. E. (2004) UCSF Chimera—a visualization system for exploratory research and analysis. Journal of computational chemistry 25, 1605–1612

109. Lourenço, E. V., and Roque-Barreira, M.-C. (2009) Immunoenzymatic quantitative analysis of antigens expressed on the cell surface (cell-ELISA) In Immunocytochemical Methods and Protocols, Springer, 301–309

110. Newton, C. L., Anderson, R. C., Katz, A. A., and Millar, R. P. (2016) Loss-of-function mutations in the human luteinizing hormone receptor predominantly cause intracellular retention. Endocrinology 157, 4364–4377

111. Zhuang, Y., Xu, P., Mao, C., Wang, L., Krumm, B., Zhou, X. E. et al. (2021) Structural insights into the human D1 and D2 dopamine receptor signaling complexes. Cell 184, 931–942. e918

